# nMAT3 is an essential maturase splicing factor required for holo-complex I biogenesis and embryo-development in *Arabidopsis thaliana* plants

**DOI:** 10.1101/2020.10.20.346734

**Authors:** Sofia Shevtsov-Tal, Corinne Best, Roei Matan, Sam Aldrin Chandran, Gregory G. Brown, Oren Ostersetzer-Biran

**Affiliations:** Department of Plant and Environmental Sciences, The Alexander Silberman Institute of Life Sciences, The Hebrew University of Jerusalem, Givat-Ram, Jerusalem 91904, Israel; School of Chemical and Biotechnology, SASTRA University, Thanjavur 613 401, India; Department of Biology, McGill University, Montreal, Quebec H3A 1B1, Canada

**Keywords:** maturase, group II intron, splicing, mitochondria, embryogenesis, Arabidopsis

## Abstract

Group II introns are large catalytic RNAs that are particularly prevalent in the organelles of terrestrial plants. In angiosperm mitochondria, group II introns reside in the coding-regions of many critical genes, and their excision is essential for respiratory-mediated functions. Canonical group II introns are self-splicing and mobile genetic elements, consisting of the catalytic intron-RNA and its cognate intron-encoded endonuclease factor (*i.e.* maturase, Pfam-PF01348). Plant organellar introns are extremely degenerate, and lack many regions that are critical for splicing, including their related maturase-ORFs. The high degeneracy of plant mitochondrial introns was accompanied during evolution by the acquisition of ‘*host-acting*’ protein cofactors. These include several nuclear encoded maturases (nMATs) and various other splicing-cofactors that belong to a diverse set of RNA-binding families, *e.g.* RNA helicases (Pfam-PF00910), Mitochondrial Transcription Termination Factors (mTERF, Pfam-PF02536), Plant Organelle RNA Recognition (PORR, Pfam-PF11955), and Pentatricopeptide repeat (PPR, Pfam-PF13812) proteins. Previously, we established the roles of MatR and three nuclear-maturases, nMAT1, nMAT2, and nMAT4, in the splicing of different subsets of mitochondrial introns in Arabidopsis. The function of nMAT3 (AT5G04050) was found to be essential during early embryogenesis. Using a modified embryo-rescue method, we show that *nMAT3*-knockout plants are strongly affected in the splicing of *nad1* introns i1, i3 and i4 in Arabidopsis mitochondria. The embryo-defect phenotype is tightly associated with complex I biogenesis defects. Functional complementation of *nMAT3* restored the splicing defects and altered embryogenesis phenotypes associated with the *nmat3* mutant-line.

## Introduction

The mitochondrion serves as a hub for cellular energy metabolism in the plant cell (Millar *et al.* 2011, Schertl and Braun 2014). As descendants from an ancestral proteobacterial symbiont, in the vast majority of species mitochondria contain their own genetic system (mitogenome, mtDNA), which encodes for rRNAs and some of the organellar tRNAs and proteins (Bonen 2018, Gualberto and Newton 2017). Plants are able to coordinate their energy demands during particular growth and developmental stages by means of nucleocytoplasmic signaling (*i.e.* nuclear to mitochondria or plastids). The metabolic functions, biogenesis and maintenance of the mitochondria are controlled by complex networks of genetic interactions between the host genome and the organelles, activities which may relate to plant terrestrialization about half a billion years ago (Best *et al.* 2020). For instance, the mitochondrial ribosomes and the energy transduction machineries are assemblies of both nuclear and organellar encoded subunits, where the correct stoichiometry in the accumulation of the different subunits composing the organellar complexes is necessary for their biogenesis and functions. These processes necessitate complex mechanisms for regulating the coordination of the expression and accumulation of the different subunits encoded by the two physically remote genomes (Fuchs *et al.* 2020, Kleine and Leister 2016, Woodson and Chory 2008). However, the identity of the messenger molecules involved in these pathways is more elusive.

The mtDNAs of land plants are remarkably variable in size, sequence and structure (Bonen 2018, Gualberto and Newton 2017). These may also contain linear or circular extrachromosomal DNAs, and undergo frequent recombination events that lead to rapid changes in genome configuration, and also result in the formation of many novel open reading frames (ORFs), many of which have no assigned functions (Bonen 2018, Gualberto and Newton 2017). Some plants have experienced dramatic expansion in mitogenome size, resulting in the largest known mitochondrial genomes (*i.e.* ~11.3 Mbp) (Sloan *et al.* 2012). Others, such as the mitochondrial genome in mistletoe (*Viscum album*) undergone massive gene loss (Maclean *et al.* 2018, Petersen *et al.* 2015, Senkler *et al.* 2018). Despite the large variation in mitogenome size and gene organization, the number of mitochondrial genes is relatively conserved in the land-plant kingdom, with about 60 known genes found in different terrestrial plant species (Bonen 2018, Grewe *et al.* 2014, Gualberto and Newton 2017, Guo *et al.* 2016, Mower *et al.* 2012, Park *et al.* 2015, Sloan, *et al.* 2012). These include tRNAs, rRNAs, ribosomal proteins, various subunits of the respiratory complexes I (NADH dehydrogenase), III (cytochrome c reductase or bc1), and IV (cytochrome c oxidase), subunits of the ATP-synthase (also denoted as CV), cytochrome *c* biogenesis (CCM) factors, and at least one component of the twin-arginine protein translocation complex.

Different from their counterparts in Animalia, the mitochondrial genes in land plants are arranged in numerous polycistronic units that are physically separated by large intergenic regions, while many of the coding regions contain intron sequences that must be excised post-transcriptionally (Zmudjak and Ostersetzer-Biran 2017). The intergenic regions are postulated to harbor important regulatory regions, such as promoters, enhancer or repressor DNA sequences, which can activate or downregulate the expression of organellar genes in a tissue, developmental or environmental dependent manner. In addition to transcriptional control, post-transcriptional RNA processing seems to play a pivotal role in the regulation of land plant mtDNA expression (Best, *et al.* 2020, Bonen 2018, Colas des Francs-Small and Small 2014, Hammani and Giege 2014, Zmudjak and Ostersetzer-Biran 2017). These involve the maturation of 5’ and 3’ termini, RNA editing (typically C-to-U exchanges) (Small *et al.* 2020) and the removal of intron sequences that reside within many essential protein-coding genes (Brown *et al.* 2014, Schmitz-Linneweber *et al.* 2015). These RNA processing steps are essential for the organellar RNAs to carry out their functions in protein synthesis, and may therefore serve as key control points in plant mitochondria gene expression.

The processes that lead to the establishment of functional mRNAs in mitochondria are accomplished largely by nuclear-encoded cofactors, which may also provide a means to link organellar functions with environmental and developmental signals (Best, *et al.* 2020). Recent data show that the nuclear genomes of angiosperms encode numerous RNA binding proteins essential for mitochondria biology and plant physiology (Colas des Francs-Small and Small 2014, Zmudjak and Ostersetzer-Biran 2017). In fact, these factors comprise a staggering proportion of the total proteome of angiosperm mitochondria, representing about 15% of the identified protein species (Fuchs, *et al.* 2020), further signifying the importance of RNA metabolism for mitochondrial biogenesis and plant physiology (Best, *et al.* 2020).

Our work focuses of group II intron splicing in land plant mitochondria. Although group II introns in plants have evolved from maturase-encoding group II sequences (Ahlert *et al.* 2006), only the fourth intron in NADH dehydrogenase 1 (*nad1* i4) in angiosperms has retained an ORF encoding a protein with sequence similarity to maturases (MatR) (Wahleithner *et al.* 1990). Previously we demonstrated that MatR is associated with *nad1* i4 and several other intron-containing pre-mRNAs, and facilitates the splicing of the introns with whom it’s associated in vivo (Sultan *et al.* 2016). In addition to MatR, genetic screens have led to the identification of many splicing cofactors in plant mitochondria (see *e.g.* (Colas des Francs-Small and Small 2014). These belong to a diverse set of RNA-binding families, *e.g.* RNA helicases (Pfam-PF00910), Mitochondrial Transcription Termination Factors (mTERF, Pfam-PF02536), Plant Organelle RNA Recognition (PORR, Pfam-PF11955), and Pentatricopeptide repeat (PPR, Pfam-PF13812) proteins, while several others are closely related to type-II intron maturases (Pfam-PF01348). For recent reviews about plant mitochondria group II intron splicing see *e.g.* (Brown, *et al.* 2014, Colas des Francs-Small and Small 2014, Hammani and Giege 2014, Schmitz-Linneweber, *et al.* 2015, Zmudjak and Ostersetzer-Biran 2017).

Arabidopsis harbours four maturase-related genes that are found in the nucleus as lone ORFs, with no trace of their corresponding group II intron sequences (Brown, *et al.* 2014, Mohr and Lambowitz 2003, Schmitz-Linneweber, *et al.* 2015). Localization analyses indicated that the four nMATs are all localized to the mitochondria, in vivo. Thus, a reasonable hypothesis is that these proteins act in the processing of mitochondrial introns in angiosperms. Previously, we established the roles of nMAT1 (At1g30010) (Keren *et al.* 2012, Nakagawa and Sakurai 2006), nMAT2 (At5g46920) (Keren *et al.* 2009, Zmudjak *et al.* 2017) and nMAT4 (Cohen *et al.* 2014) in the splicing of different subsets of mitochondrial introns in Arabidopsis.

In this work, we analyzed the RNA profiles and organellar activities associated with nMAT3 (At5g04050), and establish a role for this protein in the splicing of several mitochondrial *nad1* introns. The effects of lowering the expression of nMAT3, encoded by the At5g04050 gene-locus, on the phenotype and physiology of knockout (T-DNA) rescued lines in Arabidopsis plants are discussed.

## Results

### The Arabidopsis *nMAT3* (At5g04050) gene-locus encodes a type-II self-standing maturase factor that is expressed during early developmental stages

Group II introns are both catalytic RNAs and retroelements, which are able to transpose into homologous loci in the host genome, through a process catalyzed by a ribonucleoprotein complex consisting of the released intron and its related maturase ORF (Lambowitz and Zimmerly 2011, Zimmerly and Semper 2015). Typical maturases (MATs) are characterized by two functional domains that are required for both splicing and RNA mobility activities. These include a region with sequence similarity to retroviral-type reverse transcriptases (Pfam-PF07727), with conserved sequence blocks that are present in the fingers and palm (fingers-palm) regions and a sequence motif similar to the thumb domain of retroviral RTs (also denoted as domain X), which are associated with RNA binding and splicing (Mohr *et al.* 1993). Some, but not all, of the MATs harbor a canonical C-terminal region with homology to DNA-binding endonucleases (*i.e.* D-En domain) (Pfam-12705).

The nuclear genome of Arabidopsis encodes four maturase-related genes (annotated as nMAT’s 1 to 4) that their products are postulated to reside in the mitochondria (Brown, *et al.* 2014, Mohr and Lambowitz 2003, Schmitz-Linneweber, *et al.* 2015). Figure 1 represents a schematic representation of plant mitochondrial maturase-related proteins. MatR encodes a degenerated MAT protein that harbours some of the finger-palm regions and an intact domain-X, but lacks the C-terminal D-En domain (Sultan, *et al.* 2016). Based on their topology and predicted evolutionary origins the four nMATs in plants are classified as either type-I (*i.e.* nMAT1 and nMAT2), which harbor the RT and X domains but lack the D-En motifs, and type-II maturases (nMAT3 and nMAT4), that have retained the two domains characteristic of canonical group II intron-encoded maturases (Keren, *et al.* 2009, Mohr and Lambowitz 2003). While the RT domains in nMAT1 and nMAT2 appear to have degenerated, the D-En regions of nMAT3 and nMAT4 contain amino acid alterations that are expected to inactivate their endonuclease activities (Fig. 1a) (Mohr and Lambowitz 2003). It is therefore expected that the maturases in plants have retained their splicing activities (Cohen, *et al.* 2014, Keren, *et al.* 2009, Keren, *et al.* 2012, Nakagawa and Sakurai 2006, Zmudjak, *et al.* 2017), but lack the mobility functions associated with the model type-II group II intron-encoded endonucleases.

**Figure 1.**
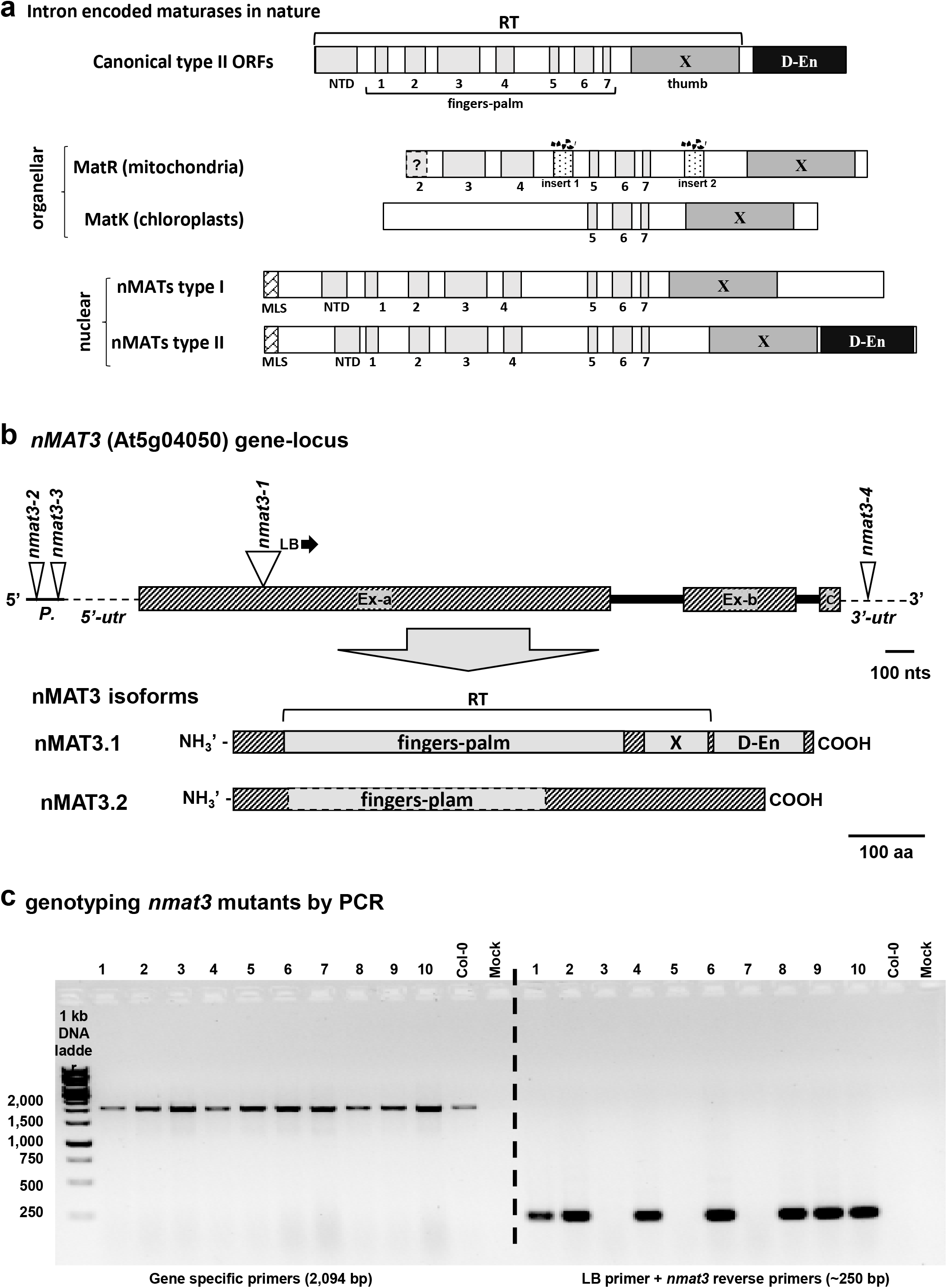
Schematic representation of the *nMAT3* (At5g04050) gene. (a) Schematic representation of plant mitochondrial maturase-related proteins. The figure is modified from (Sultan, *et al.* 2016). Shaded boxes represent different motifs associated with model group II intron-encoded endonucleases. These include the reverse transcriptase (RT) domain (Pfam-PF01348), with its intrinsic NTD (N-terminal domain), the finger-palm (RT I to VII motifs), and the RNA-binding and splicing subdomains (*i.e.* domain X or thumb; Pfam-08388). Some members of this family also harbor an additional C-terminal DNA binding and endonuclease domain (D-En, Pfam-12705). (b) Schematic representation of the *nMAT3* gene and its two related gene-products, nMAT3.1 and nMAT3.2. The different motifs are highlighted within the sequence of *nMAT4*, according to Mohr and Lambowitz (2003) and Sultan, *et al.* (2016). The location of the T-DNA insertion sites is indicated. (c) PCR-based genotyping of the progeny from heterozygous *nmat3* (SAIL_254_F03) plants. Primers are listed in Table S4.

According to ‘The Arabidopsis Information Resource’ (TAIR; http://www.arabidopsis.org), the At5g04050 gene-locus (encoding the nMAT3 protein) (Keren, *et al.* 2009, Mohr and Lambowitz 2003), harbors two intron sequences that are suggested to be alternatively spliced into two isoforms: a 3,027 nucleotide-long unspliced product (At5g04050.1), encoding a 599 amino acids protein (*i.e.* nMAT3.1), and a spliced (At5g04050.2) mRNA variant of 2,905 nucleotides, encoding a 694 amino acid long protein (nMAT3.2) (Fig. 1b and Supplemental Figure S1). Yet, DNA sequencing of the At5g04050 gene-locus revealed to some errors within the sequence of *nMAT3* intron 1 (Fig. S1). BLAST searches indicated that the modified intron sequence is conserved in different Brassicales species, *e.g. Arabis alpine*, *Arabidopsis lyrate*, *Brassica rapa*, *Brassica napus*, *Brassica oleracea*, *Camelina sativa* and *Raphanus sativus* (Fig. S2). The putative open-reading-frame of the modified At5g04050 gene-locus encodes a 757 amino acids product (instead of 599 aa’s) for the unspliced *nMAT3.1* isoform. Analyses of nMAT3.1 and nMAT3.2 by the SMART (Letunic *et al.* 2012) and Conserved Domain Database (CDD) (Marchler-Bauer *et al.* 2003) servers indicated that the nMAT3.2 variant harbors some of the fingers-palm regions, but lacks the X and D-En domains, whereas the ‘unspliced’ nMAT3.1 isoform harbors an both the RT and D-En motifs, typical to group II intron-encoded maturases (Fig. 1b). The expression of the two putative At5g04050 gene products was further analyzed by RT-PCR, with primers flanking introns 1 and 2. These analyses indicated the existence of only a single isoform, the ‘unspliced’ *nMAT3.1* gene-product in *Arabidopsis thaliana* (var. Col-0) plants (Fig. S3). Based on these data it is predicted that Arabidopsis harbors only a single functional isoform, the *nMAT3.1* gene-product, which encodes an ‘intact’ maturase-related (nMAT3) protein.

### *nMAT3* encodes a lowly-expressed protein that is localized to the mitochondria, and have essential roles during early embryo-development

The TAIR and the ‘Genevestigator analysis toolbox’ (Hruz *et al.* 2008, Zimmermann *et al.* 2004) servers indicate that At5g04050 is a lowly expressed gene, which shows a differential expression throughout development in different organs, with dominant expression in embryonic organs, and young developing tissues, *e.g.* leaves, apical root meristem, flowers and the shoot apex (Fig. S4). The gene expression databases indicate that nMAT3 expression is mostly upregulated during germination and early plant life (Fig. S4). In accordance with its predicted low-expression levels, nMAT3 could not be found among the different proteins identified in mass-spectrometry analyses of plant organellar fractions, which contain various RNA binding cofactors (*i.e.* SUBA4 server) (Hooper *et al.* 2017). nMAT3 is predicted to reside in the mitochondria by the consensus SUBAcon (Hooper, *et al.* 2017). Previously, we demonstrated by GFP localization analyses that the Arabidopsis *nMAT3* gene-locus encodes a mitochondria-localized protein (Keren, *et al.* 2009). The homology of At5g04050 gene product(s) with model group II-encoded maturases and its localization by GFP-targeting studies (Keren, *et al.* 2009) support a role for nMAT3 in the processing of organellar introns in plants.

To study the putative roles of nMAT3 in mitochondria biogenesis, we examined available T-DNA insertional lines at the At5g04050 gene-locus, available at TAIR’s seed stock. These include heterozygous SAIL_254_F03 (*nmat3-1*) line, containing a T-DNA insertion within the RT domain of nMAT3, the homozygous SALK_011307C (*nmat3-2*) and heterozygous SALK_139392 (*nmat3-3*) lines that contain an insertion within the putative promoter region of At5g04050 gene, and heterozygous SALK_144082 (*nmat3-4*) line that contains a T-DNA insertion that is mapped to the 3’-UTR in *nMAT3* gene. Sequencing of genomic PCR products, spanning the T-DNA insertional junctions in the four *nmat3* mutants confirmed the integrity of the insertional lines (Fig. 1b, and Fig. S1). No plants homozygous for the *nMAT3-1* mutant allele could be recovered among the seeds obtained from heterozygote SAIL_254_F03 line (Fig. 1c). Homozygous *nmat3-2*, *nmat3-3* and *nmat3-4* plants (containing T-DNA insertions at the 5’ or 3’ termini of the *nMAT3* gene) did not show any obvious phenotypes under normal growth conditions (Keren, *et al.* 2009). RT-qPCR analysis of nMAT3 expression indicated that the T-DNA insertions did not alter the expression of nMAT3 in the homozygous lines (Keren, *et al.* 2009).

The selfed progeny of the heterozygous *nmat3-1* mutants showed that about a quarter of the seeds did not germinate (*i.e.* 28 out of 101 seeds), suggesting that *nMAT3* encodes an essential gene. The heterozygous *nmat3-1* plants were phenotypically indistinguishable from the wild-type plants (Col-0), suggesting that the insertion within the coding region of *nMAT3* gene result in a recessive embryo-lethal phenotype, and that a single copy of the gene is sufficient to support normal growth and development. Microscopic analysis of siliques obtained from Arabidopsis wild-type (Col-0; Fig. 2a) and heterozygous *nmat3-1* plants (Fig. 2b) showed the presence of translucent seeds in *nmat3-1* mutant-line (Fig. 2b, marked with black arrow). These comprised 24.6 % of 693 seeds obtained from 43 siliques of *nmat3-1* plants, and 0% white seeds out of 160 seeds obtained from wild-type plants. While seeds were produced normally in Col-0 (Fig. 2c), the mature siliques of *nmat3-1* contained about 25% darkened and shrunken seeds (Fig. 2d). Nomarski microscopy analyses further indicated that green seeds carry mature embryos, while the translucent seeds of the same progeny contain embryos arrested at the heart-torpedo transition stage (Fig. 2e-f). During the maturation of the siliques, some of the pale seeds collapsed into seeds lacking embryos (Fig. 2e, indicated by an arrow), while others degenerate into shrunken-brown seeds (Fig. 2e, indicated by an arrow). The morphology of the shrunken seeds resembles that of *wrinkled1* mutants (Focks and Benning 1998), a seed developmental defect that is also associated with altered mitochondrial activities (Keren, *et al.* 2012, Kuhn *et al.* 2015, Pinfield-Wells *et al.* 2005). While wild-type (Fig. 2g) and ‘normal looking’ seeds collected from heterozygous *nmat3-1* plants germinate successfully on soil or MS-agar plates (Fig. 2g), wrinkled seeds from the same siliques fail to sprout, and did not germinate even when incubated on (1%) sucrose-containing MS-agar medium (Fig. 2d), a condition used to recover germination-arrested phenotypes in Arabidopsis mutants affected in mitochondria biogenesis, e.g., the *cod1* (Dahan *et al.* 2014) and *cal1/cal2* mutants (Cordoba *et al.* 2016, Fromm *et al.* 2016a).

**Figure 2.**
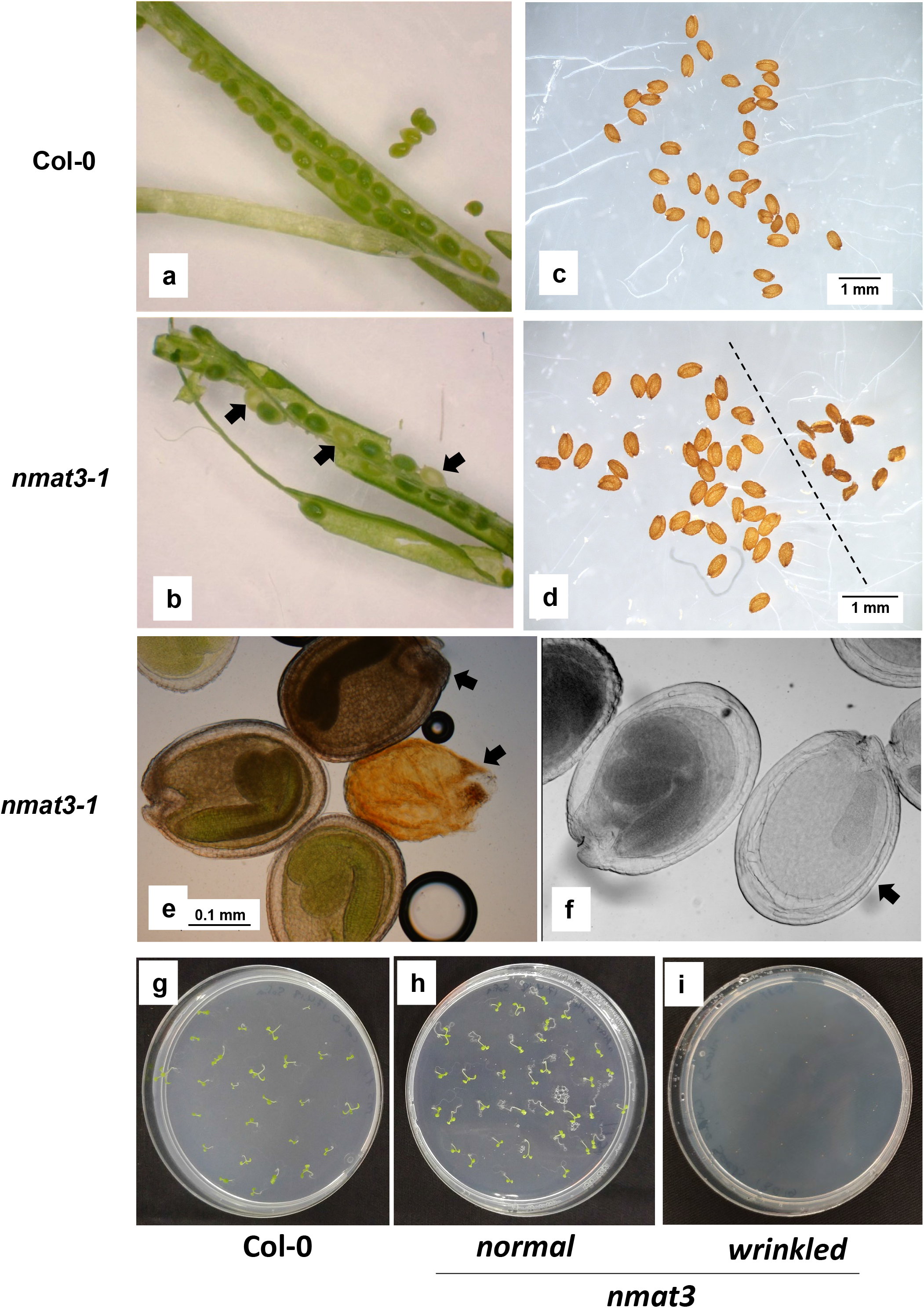
Plant phenotypes associated with *nmat3* mutations. Mature siliques were obtained from Arabidopsis wild-type (Col-0) and heterozygous *nmat3-1* plants. The effects of loss of nMAT3 on seed development and morphology (a-f), and seed germination (g-i) of Arabidopsis wild-type (Col-0) (panels a, c, and g) and knockout *nmat3-1* mutant line (b-f). Embryo-developmental phenotypes in green and white seeds of the same progeny of *nmat3-1* plants. Images taken with differential interference contrast (*i.e.* Nomarski) microscopy. Bars in panels c and represent 1 mm and 0.1 mm in panels e and f.

### Embryo rescue of Arabidopsis *nMAT3-1* mutant-line

Arabidopsis mutants impaired in mitochondria biogenesis or functions often display altered seed germination, a phenotype that is associated with embryo developmental defects (Colas des Francs-Small and Small 2014, Cordoba, *et al.* 2016, Dahan, *et al.* 2014, Focks and Benning 1998, Franzmann *et al.* 1989, Fromm, *et al.* 2016a, Keren, *et al.* 2009, Kuhn, *et al.* 2015, Ostersetzer-Biran 2016, Pinfield-Wells, *et al.* 2005). Our data show that loss-of-function knockout *nMAT3-1* mutant-seeds were not informative for the analysis of nMAT3 functions in Arabidopsis plants. None of the mature wrinkled seeds of *nmat3-1*, which we considered to be homozygous for the *nMAT3-1* allele, were able to germinate. One approach for studying the *post*-embryonic function of essential genes involves a partial complementation of the mutation, by cloning a target gene under a seed-specific promoter, *e.g. ABI3* (Despres *et al.* 2001) or *RPS5A* (Johnson *et al.* 2008) promoters. The technique has been successful in studying various embryo arrested mutants in Arabidopsis (Despres, *et al.* 2001), among which some are affected in organellar RNA metabolism (Aryamanesh *et al.* 2017, Sun *et al.* 2018). However, this method was found unsuitable for the analysis of the *nmat3-1* mutant, as no individuals homozygous for the *nMAT3-1* allele could be recovered among the progeny of heterozygous *nmat3-1* plants carrying the *ABI:nMAT3* transgene.

In addition to the use of seed-specific promoters, embryo-rescue approaches have been previously used to recover germination-arrested phenotypes in plant mutants affected in early embryogenesis (Franzmann, *et al.* 1989), some of which show altered mitochondria biogenesis and function, *e.g. cod1* (Dahan, *et al.* 2014), *ndufv1* (Kuhn, *et al.* 2015) and *cal1/cal2* mutants (Cordoba, *et al.* 2016, Fromm, *et al.* 2016a). Unfortunately, the settings described for the rescue of *cod1*, *ndufv1* or *cal1/cal2* mutants, were found insufficient to allow the germination of *nmat3-1* seeds, which may relate to the early embryogenesis defect phenotype of *nmat3-1* mutant (Fig. 3), whereas the seeds of *cod1* and *cal1/cal2* seem to contain small, but otherwise to fully developed embryos (Cordoba, *et al.* 2016, Dahan, *et al.* 2014, Fromm, *et al.* 2016a).

**Figure 3.**
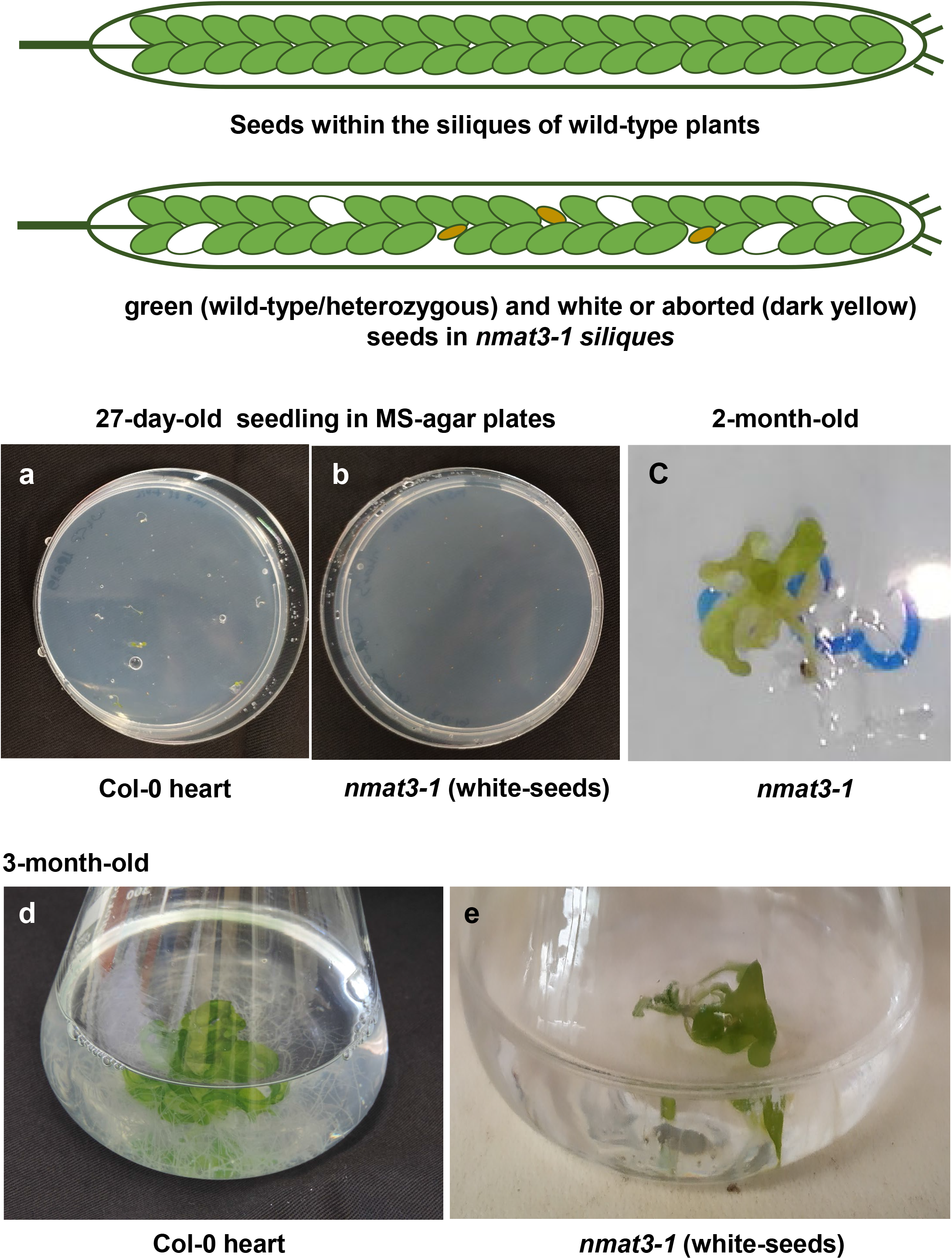
Plant phenotypes associated with rescued homozygous *nmat3-1* mutant line. Seeds were collected under sterile conditions from surface sterilized immature siliques of Col-0 and heterozygous *nmat3-1* plants. While the seeds of wild-type plants show normal development, the siliques of *nmat3-1* mutants contain green (*i.e.* wild-type or heterozygous embryos), white (homozygous *nmat3* embryos) and aborted (degenerated embryos) *nmat3-1* mutant seeds (dark yellow). Pictures represent embryo rescue of (**a**) wild-type seeds at the heart stage and (**b**) white seeds collected from the immature siliques of heterozygous *nmat3-1* plants, following 27 days of incubation on MS-agar plates supplemented with vitamins and sugars (see Experimental procedures). (**c**) A 2-month-old rescued homozygous nmat3-1 seedling at the L6 stage. (**d**) 3-month-old rescued wild-type plants transferred to liquid MS-media supplemented with 1% sucrose and vitamins (see Experimental procedure). (**e**) Image of 3-month-old rescued *nmat3-1* plantlet in the liquid MS-media.

We tried to overcome this problem, by assaying the germination of *nmat3-1* seeds collected from green-matured siliques (*i.e.* about 10 days post anthesis) (Mizzotti *et al.* 2018), using various growth media and growth conditions. The optimized settings for germination of homozygous *nmat3-1* were obtained when white-translucent seeds, selected from sterilized immature siliques of heterozygous plants, were sown on MS-agar plates supplemented with 1% sucrose and vitamins (Fig. 3), conditions that we found optimal for the establishment of a PPR-related (i.e. *msp1025*) mutant, which is also arrested at the late heart stage (Best *et al.* 2019). Under the in vitro conditions, many seeds of wild-type plants containing embryos at the late heart or torpedo stages (Fig. 3a) and a few of the cultivated *nmat3-1* seeds germinated after 4~5 weeks (Fig. 3b), whereas a total of about 90% of green seeds and 40% of the white seeds germinated during 3 months of incubation. Importantly, genotyping by PCR indicated that seedlings obtained from white-translucent seeds were also homozygous for the loss-of-function *nMAT3* allele. These slowly developed to the six-true-leaf stage (Boyes *et al.* 2001) following 2 months of incubation in the growth chamber, but were unable to continue their development beyond the L6-stage (Fig. 3c). When transplanted into a liquid MS-based medium containing 1% sucrose and vitamins (see Experimental procedures) their development proceeded beyond the L6 stage, and a significant increase in leaf and root biomass was apparent in the culture medium following 4 weeks (Fig. 3d). None of the rescued *nmat3* plants were able to complete a life cycle and produce viable seeds. For the sequential analyses of the RNA and protein profiles of *nmat3-1* mutants, we used 3 to 4-month-old embryo-rescued plantlets obtained from white seeds obtained from the siliques of heterozygous *nmat3* mutant plants. As control we used embryo-rescued wild-type (Col-0) seedlings, collected from seeds containing embryos at the late heart stage. 3-week-old Arabidopsis MS-grown seedlings were used as a reference for the molecular and biochemical assays used in this study.

### Mutations in the *nMAT3* gene-locus strongly affect the accumulation or processing of mature *nad1* transcripts in Arabidopsis mitochondria

To gain more insight into the putative roles of nMAT3 in mtRNA metabolism, we used transcriptome analyses of embryo-rescued wild-type and *nmat3-1* plants by quantitative reverse transcription PCR (RT-qPCR) (Cohen, *et al.* 2014, Colas des Francs-Small *et al.* 2012, Keren, *et al.* 2012, Sultan, *et al.* 2016, Zmudjak *et al.* 2013). The relative accumulation of mitochondrial transcripts (mtRNAs) was estimated by RT-qPCR from the ratios between embryo-rescued *nmat3-1* plantlets versus those of 3-week-old MS-agar grown wild-type plants (Fig. 4a). Several genes in Arabidopsis mitochondria are interrupted by group II introns, i.e., *ccmFc*, *cox2*, *nad1*, *nad2*, *nad4*, *nad5*, *nad7*, *rpl2* and *rps3* (Unseld *et al.* 1997). Moreover, several introns have been disrupted by DNA rearrangements, so that separately-transcribed precursors undergo splicing in ‘*trans*’. Accordingly, the maturation of *nad1* involves the joining of five exons, encoded by three individually-expressed gene fragments (*i.e*. *nad1.1*, *nad1.2*and *nad1.3*, which are spliced through two *trans*- (introns 1 and 3) and two *cis*- (introns 2 and 4) events. Likewise, the maturation of *nad2* involves the splicing of 3 *cis*- (*i.e. nad2* introns 1, 3 and 4) and one *trans*-spliced intron (*nad2* i2) found on two individual transcripts (*nad2.1* and *nad2.2*), whereas the maturation of *nad5* involves the joining of five exons, encoded by three individually-expressed gene fragments (*i.e*. *nad5.1*, *nad5.2* and *nad5.1*), where introns 2 and 3 are physically separated on the mitogenome and are spliced in ‘*trans’*.

**Figure 4.**
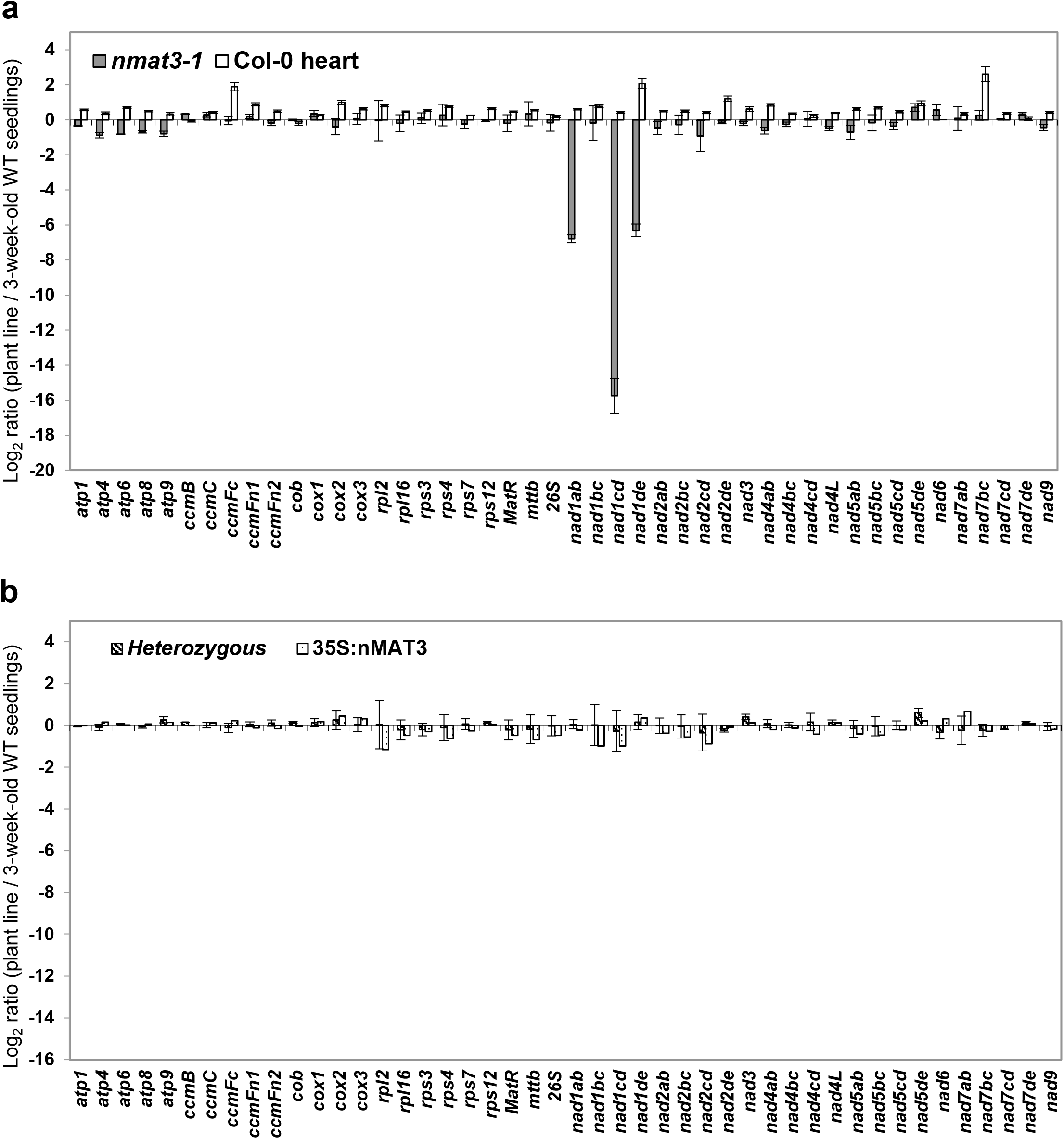
Abundance of mitochondrial transcripts in the *nmat3-1* mutant. Transcriptome analyses of mitochondrial mRNAs levels in Arabidopsis plants by RT-qPCR was performed essentially as described previously (Cohen, *et al.* 2014, Sultan, *et al.* 2016, Zmudjak, *et al.* 2013). RNA extracted from 3-week-old seedlings of wild-type plants, 4~5 months old rescued *nmat3-1* mutant plants and plantlets obtained from the immature seeds of Col-0 plants containing embryos at the heart stage (Fig. 3), was reverse-transcribed, and the relative steady-state levels of cDNAs corresponding to the different organellar transcripts were evaluated by qPCR with primers which specifically amplified mRNAs (Supplemental Table S1a). Histograms showing the relative mRNAs levels (*i.e*. log2 ratios) in (**a**) rescued homozygous *nmat3-1* mutant and rescued Col-0 plantlets, or (**b**) heterozygous *nmat3-1* mutants and *nmat3:35S-nMAT3* complemented plants, versus those of 3-week-old MS-grown wild-type plants. Transcripts analyzed in these assays include the NADH dehydrogenase (CI) subunits *nad1* exons a-b, b-c, c-d, d-e, *nad2* exons a-b, b-c, c-d, d-e, *nad3*, *nad4* exons a-b, b-c, c-d, *nad4L*, *nad5* exons a-b, b-c, c-d, d-e, *nad6* subunit, *nad7* exons a-b, b-c, c-d, d-e, and *nad9*, the complex III *cob* subunit, cytochrome oxidase (complex IV) *cox1*, *cox2* exons a-b and *cox3* subunits, the ATP synthase (i.e., complex V) subunits *atp1*, *atp4*, *atp6*, *atp8* and *atp9*, genes encoding the cytochrome c maturation (*ccm*) factors, *ccmB*, *ccmC*, *ccmFn1*, *ccmFn2* and *ccmFc* exons a-b, the ribosomal subunits *rpl2* exons a-b, *rps3* exons a-b, *rps4*, *rps7*, *rps12*, *rpl16*, *rrn26*, and the *mttB* gene. The values are means of nine RT-qPCR reactions corresponding to three biological replicates (error bars indicate one standard deviation), after normalization to the *GAPDH* (AT1G13440)*, ACTIN2* (At3g1878), *18S-rRNA* (At3g41768), *RRN26* (*i.e.* mitochondrial 26S-rRNA, Atmg00020), *RRN5* (Atmg01380) and *RRN18* (Atmg01390) genes.

The accumulation of the entire set of splice variants was examined by RT-qPCR with oligos designed to different exon-exon and exon-intron or intron-exon pairs (Table S1). As shown in Figure 4a, notable reductions in mtRNA were apparent for different *nad1* transcripts in *nmat3-1* plantlets, where the steady-state levels of transcripts corresponding to *nad1* exons a-b (*i.e. nad1ab*) was about 110-fold lower than that of the wild-type plants and about 57,000-fold lower in the case of *nad1cd* transcript, while the accumulation of *nad1de* transcript was about 80-fold-lower than that of wild-type plants. The expression of other group II intron-containing transcripts, including *ccmFc*, *cox2*, *nad4*, *nad5*, *nad7*, *rpl2 rps3*, was not significantly affected in the homozygous *nmat3-1* mutant plants. Likewise, no significant effects on the accumulation of mtRNAs corresponding to ‘intronless’ genes were apparent in the mutant (Fig. 4a). These included *cox1* and *cox3* subunits of complex IV, subunits of the ATP synthase complex (CV), cytochrome C biogenesis and maturation (*ccm*) factors, and various ribosomal genes, which their mRNAs levels in *nmat3-1* were generally comparable to those seen in wild-type plants (Fig. 4a). The roles of nMAT3 in the maturation or processing of *nad1* transcripts, was also indicated by northern blot analyses (Fig. S5). Reduced *nad1* mRNA was also found for *nmat1* mutants affected in the splicing of introns found in *nad1*, *nad2* and *nad4* (Keren, *et al.* 2012, Nakagawa and Sakurai 2006), while reduced *nad1*, *nad4*, *nad7* and *cox2* mRNA levels are found in in *nmat2* mutants (Keren, *et al.* 2009, Zmudjak, *et al.* 2017) coincide with splicing defects in many of the organellar introns (Fig. S5).

Here, we also analyzed the mtRNA profiles of rescued Col-0 plantlets (Fig. 4a, Col-0 heart), to analyze changes in mtRNA metabolism due to tissue-specific or or developmental effects, as observed during the germination of wheat seeds (Li-Pook-Than *et al.* 2004). The RNA profile of rescued wild-type embryos at the heart stage was similar to that of the 3-week-old MS-grown wild-type plants, though the steady-state levels of many of the transcripts were slightly upregulated (about 1.5x ~ 4.0x) in the rescued plantlets (Fig. 4a). No significant changes in the accumulation of mtRNAs was evident in heterozygous *nmat3-1* (Fig. 4b). We thus concluded that the observed defects in the expression or processing of *nad1* transcripts in *nmat3-1* relate to RNA metabolism defects and not to developmental effects between embryo-rescued seedlings and 3-week-old MS-grown plants.

### nMAT3 is required for the splicing of *nad1* introns 1, 3 and 4 in Arabidopsis mitochondria

The four maturases in Arabidopsis, including the mitochondria-encoded MatR factor (Sultan, *et al.* 2016), as well as the nuclearly encoded nMAT1 (Keren, *et al.* 2012, Nakagawa and Sakurai 2006) and nMAT2 (Keren, *et al.* 2009, Zmudjak, *et al.* 2017) and nMAT4 (Cohen, *et al.* 2014) protein, are all function in the splicing of group II introns in Arabidopsis mitochondria. To examine potential effects by nMAT3 on the splicing of *nad1* introns 1, 3 and 4, we compared the splicing efficiencies (*i.e*. the ratios of pre-RNAs to mRNAs in mutants versus wild-type plants) of the 23 introns in Arabidopsis mitochondria (Unseld, *et al.* 1997), between wild-type and *nmat3*-1 plants (Fig. 5). Splicing defects were determined to be present by the RT-qPCR analyses in cases where the accumulation of a specific pre-RNA was correlated with a reduced level of its corresponding mRNA in the mutant. The analyses indicated strong defects in the maturation of *nad1* exons a-b, c-d and d-e, where the splicing efficiencies for the splicing of *nad1* intron 1 (*nad1* i1), *nad1* i3 and *nad1* i4 were reduced by about 7-fold, 19,000-fold and 352-fold, respectively, in comparison to those of wild-type plants (Fig. 5a). Reduced splicing efficiencies in *nmat3-1* were also noted for *nad2* i1 (about 5.5-fold) and *nad2* i2 (about 2.2-fold) (Fig. 5a). The splicing efficiencies of the 23 mitochondrial introns of rescued wild-type embryos (Col-0 heart) and heterozygous *nmat3-1* plants were equivalent to those of the 3-week-old MS-gown wild-type plants (Fig. 5b). These data indicate that nMAT3 is pivotal for the *trans*-splicing of *nad1* i3, but is also required for the efficient *cis* or *trans*-splicing of *nad1* introns 1 (*trans*) and 4 (*cis*) and *nad2* introns 1 (*cis*) and 2 (*trans*).

**Figure 5.**
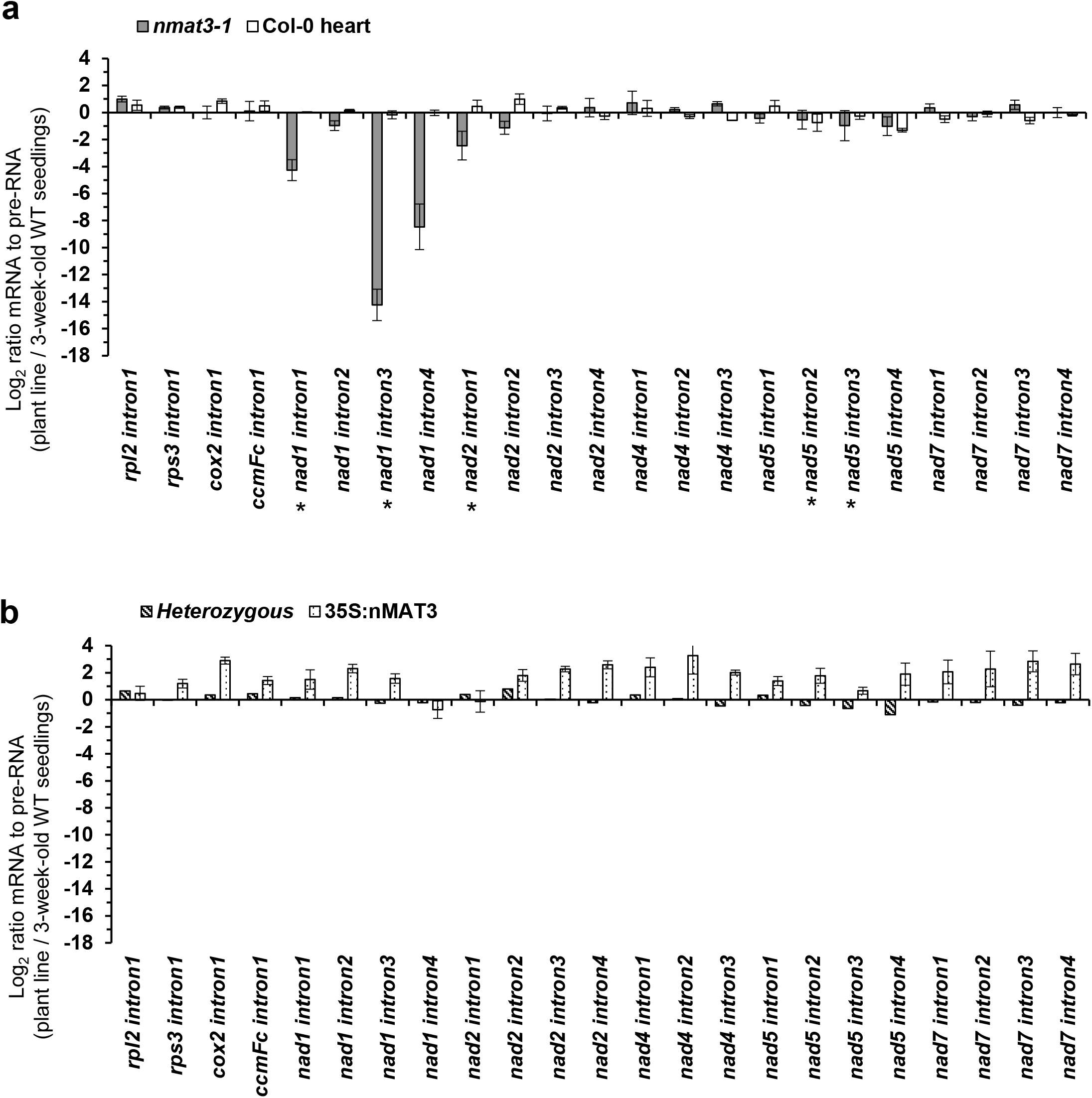
Splicing efficiencies in *nmat3-1* mutants. The relative accumulation of mRNA and pre-RNA transcripts in wild-type and rescued homozygous *namt3-1* plants, corresponding to the 23 group II intron sequences in Arabidopsis, was evaluated by RT-qPCR, essentially as described previously (Cohen, *et al.* 2014, Sultan, *et al.* 2016). RNA extracted from wild-type (Col-0), embryo-rescued *nmat3* mutants or Col-0 plants at the heart stage (Fig. 3) and *nmat3:35S-nMAT3* complemented plants was reverse-transcribed, and the relative steady-state levels of cDNAs corresponding to the different organellar transcripts were evaluated by qPCR with primers which specifically amplified pre-RNAs and mRNAs (Supplemental Table S1b). The histograms show the splicing efficiencies as indicated by the log2 ratios of pre-RNA to mRNA transcript abundance in (**a**) rescued homozygous *nmat3-1* mutant and rescued Col-0 plantlets, or (**b**) heterozygous *nmat3-1* mutants and *nmat3:35S-nMAT3* complemented plants, versus those of 3-week-old MS-grown wild-type plants. The values are means of 15 RT-qPCR reactions corresponding to five biological replicates (error bars indicate one standard deviation), after normalization to the *GAPDH* (AT1G13440)*, ACTIN2* (At3g1878), *18S-rRNA* (At3g41768), *RRN26* (*i.e.* mitochondrial 26S-rRNA, Atmg00020), *RRN5* (Atmg01380) and *RRN18* (Atmg01390) genes.

### Rescuing *nmat3-1* the growth phenotypes and mtRNA metabolism by a constitutive expression of the nMAT3 gene

To confirm that the observed growth phenotypes and altered organellar RNA metabolism are indeed associated with the loss of nMAT3, we transformed the At5g04050 gene to heterozygous *nmat3-1* plants to facilitate a ‘functional complementation’ of the mutation. For this purpose, the Arabidopsis *nMAT3* gene was obtained by PCR, cloned into the shuttle vector pCAMBIA-2300 under the control of the constitutive 35S-promoter, and transformed to heterozygous *nmat3-1* mutants by the floral-dip method (see Experimental procedure). A single transformant line, which contained the *35S:nMAT3* transgene in the background of a homozygous *nmat3-1* mutant-line, was recovered among the progeny of T2 transformed plants (*i.e. nmat3*:35S nMAT3). Seeds obtained from the complemented line were germinated on MS-agar medium, and 3-week-old seedlings of *nmat3*:35S-nMAT3 seedlings were analyzed in regard to their associated organellar RNA profiles. The RT-qPCR analyses indicated that the reduced accumulation of *nad1* and *nad2* transcripts (Fig. 4b), as well as the splicing defects we see in *nad1* introns 1,3 and 4 and *nad2* intron 1 and 2 (Fig. 5b) were fully restored in *nmat3*:35S-nMAT3 plants. The genetic data provide with a strong indication that the mutation in *nMAT3* gene-locus is directly associated to the mtRNA defects seen in *nmat3-1* mutant-line.

### Analysis of the biogenesis of organellar respiratory chain complexes in *nmat3* mutants

The respiratory machinery is made of four major electron transport carriers, denoted as complexes I to IV (*i.e.* CI, CII, CIII and CIV) and the ATP synthase enzyme (CV), as well as various proteins involved in non-phosphorylating bypasses of the electron transport chain (ETC), *i.e.* alternative oxidases (AOXs) and rotenone-insensitive NAD(P)H dehydrogenases (NDs) (Jacoby *et al.* 2012, Millar, *et al.* 2011, Schertl and Braun 2014, Senkler *et al.* 2017, Subrahmanian *et al.* 2016). The altered mtRNA metabolism we saw in *nmat3-1* were expected to result in major effects on mitochondrial activities and plant physiology, *e.g.* (Brown, *et al.* 2014, Colas des Francs-Small and Small 2014, Zmudjak and Ostersetzer-Biran 2017). NAD1 is essential for CI biogenesis and function (Braun *et al.* 2014, Fromm *et al.* 2016b, Ligas *et al.* 2019, Ostersetzer-Biran 2016). The defects we see in the splicing of *nad1* introns 1,3 and 4 (Figs. 4 and 5) suggest that NAD1 is absent, or exists in very low levels, in the mitochondria of *namt3-1* mutant plants. The steady-state levels of various CI subunits were examined in 3-week-old MS-grown plants, heterozygous *nmat3-1* plants, rescued Col-0 and *nmat3-1* and complemented *nmat3*:35S-nMAT3 line by immunoblot analyses with antibodies raised against CA2 (~30 kDa) (Perales *et al.* 2005, Sunderhaus *et al.* 2006), NAD1 (~36 kDa) and NAD9 (~23 kDa) (Lamattina *et al.* 1993). The immunoblot assays indicated that while the steady-state levels of NAD9 and CA2 subunits are equivalent to those of 3-week-old MS-grown wild-type plants, a protein band of *ca.* 50 kDa that corresponds to NAD1 protein was not visible in the *nmat3-1* mutant line (Fig. 6a).

**Figure 6.**
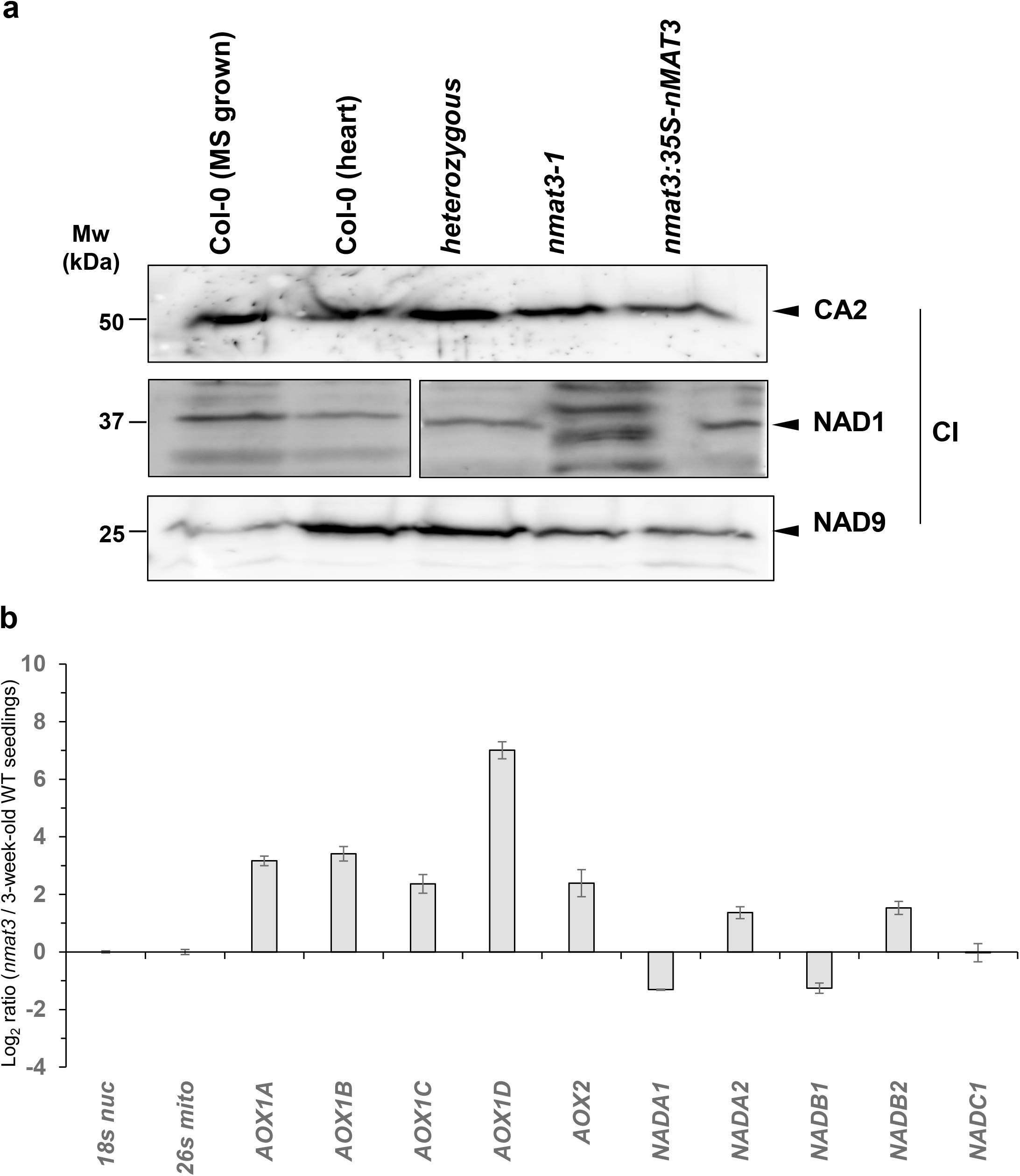
Relative accumulation of organellar proteins in wild-type and *nmat3-1* plants. (**a**) Immunoblots with total proteins (about 50 μg), extracted from 3-week-old MS-grown wild-type (Col-0) plants, rescued Col-0 (heart) or *nmat3-1* plants, and complemented *nmat3-1:35S-nMAT3* mutants. The blots were probed with polyclonal antibodies raised to NADH-oxidoreductase subunit 1 (NAD1), γ-carbonic anhydrase-like subunit 2 (CA2) and NAD9 proteins. (**b**) Analyses of the expression levels of different alternative oxidases (AOXs) and rotenone-insensitive NAD(P)H dehydrogenases (NDs) in Arabidopsis plants was performed by RT-qPCR, essentially as described previously (Cohen, *et al.* 2014, Sultan, *et al.* 2016, Zmudjak, *et al.* 2013). RNA extracted from wild-type (Col-0) and embryo-rescued *nmat3-1* mutant plants, was reverse-transcribed, and the relative steady-state levels of cDNAs corresponding to the different *AOX* and *ND* transcripts were evaluated by qPCR. The histogram shows the relative mRNAs levels (*i.e*. log2 ratios) in rescued *nmat3* mutant line and Col-0 plantlets, versus 3-week-old MS-grown wild-type plants.

The relative abundances of native respiratory complexes I, III, IV and V in 3-week-old wild-type plants, embryo-rescued *nmat3-1* and Col-0 seedlings (Col-0 heart), was analyzed by Blue-native (BN) PAGE, followed by ‘in-gel’ activity assays and immunoblot analyses (Fig. 7). Arrows indicate to the native complexes I (~1,000 kDa), CIII dimer (III_2_, ~500 kDa), the supercomplex I+III_2_ (about 1,500 kDa) (Braun, *et al.* 2014, Senkler, *et al.* 2017), CIV (~220 kDa), and CV (~600 kDa). The BN-PAGE analyses indicated that respiratory chain CI is below detectable levels in homozygous *nmat3-1* plants (Fig. 7, and Table S2). No protein bands that correspond to holo-CI (CI) or to supper complexes I_2_III_4_ or I-III_2_ (Braun, *et al.* 2014, Ligas, *et al.* 2019, Ostersetzer-Biran 2016, Sunderhaus *et al.* 2010) were observed in BN-gel activity assays and immunoblots of native gels with antibodies raised to the CA2 subunit of the membranous arm or the NAD9 subunit of the soluble arm of CI (Fig. 7). The immunoblots with CA2 subunit CI further indicated the presence of lower molecular weight bands of *ca.* 550, 500, 450 and 200 kDa in *nmat3-1* (Fig. 7, labeled as I*). Based on their appearance in the CA2 blot these particles were related to CI assembly intermediates of the membrane arm which are also apparent in various other Arabidopsis mutants affected in CI biogenesis (Braun, *et al.* 2014, Meyer 2012, Meyer *et al.* 2011, Ostersetzer-Biran 2016). In support of this assumption, the lower mass CI particles are not apparent in in the NAD9 blot (Fig. 7). No significant changes in CI accumulation (i.e., CA2 and NAD9 blots) or activity were observed in the heterozygous nmat3-1 line, nor in BN-PAGE analyses of rescued Col-0 plants (Fig. 7, and Table S2). We noticed that the activities and accumulation of CI were upregulated in embryo-rescued Col-0 plants, as indicated by the *in-gel* activity assays and immunoblots of CA2 subunit (*i.e.* 1.3-fold increase) (Fig 7, and Table S2).

**Figure 7.**
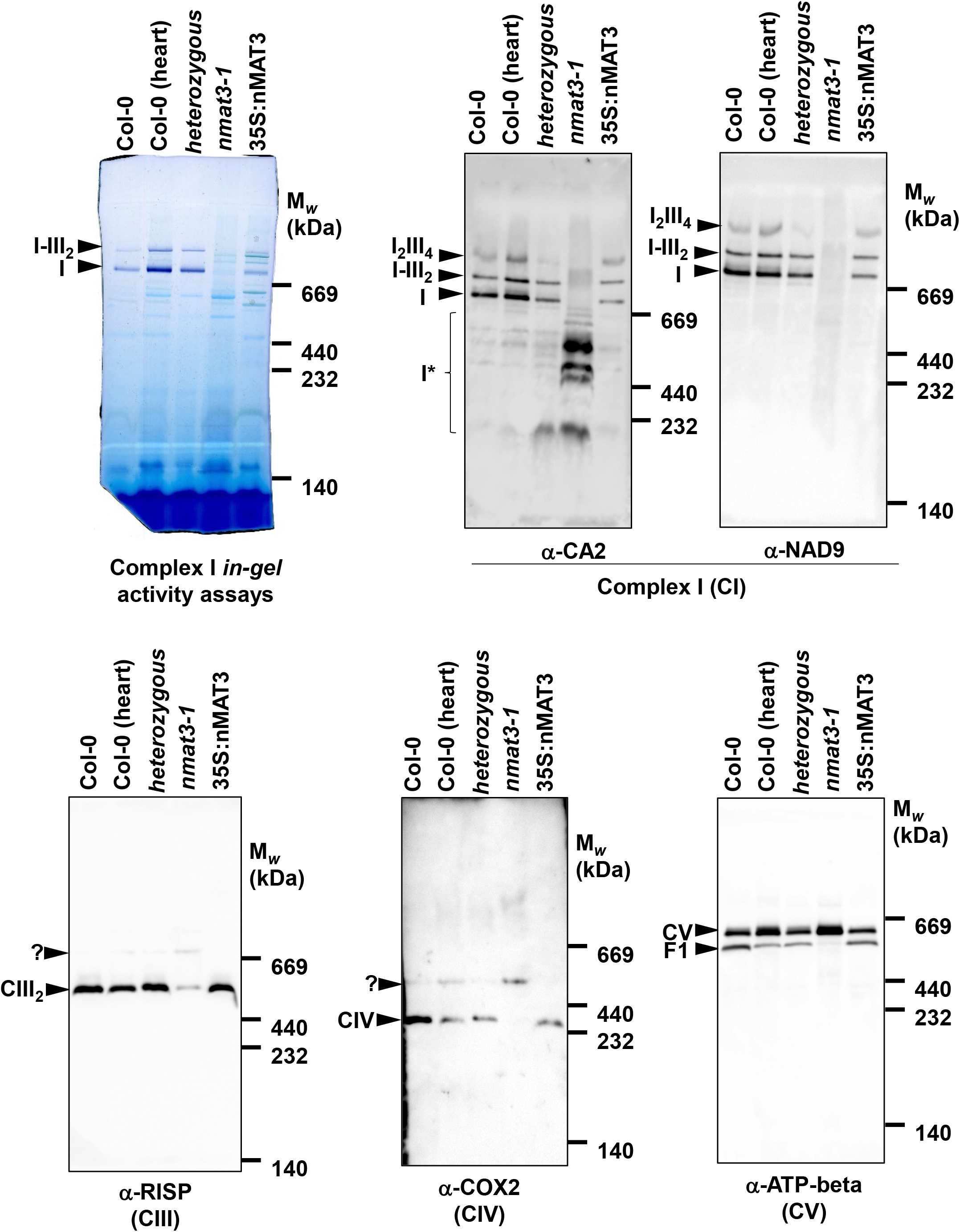
Relative accumulation of native organellar complexes in *nmat3-1* mutants. Blue native (BN)-PAGE analyses of crude organellar membranous fractions were generally performed according to the method described by Pineau, *et al.* (2008) and Sultan, *et al.* (2016). An aliquot equivalent to 40 mg of crude Arabidopsis mitochondria extracts, obtained from wild-type (Col-0), rescued Col-0 plants (heart) or *nmat3-1* plants, and complemented *nmat3-1:35S-nMAT3*, was solubilized with 5% (w/v) digitonin, and the organellar complexes were resolved by BN-PAGE. For immunodetection, the proteins were transferred from the native gels onto a PVDF membrane. The membranes probed with antibodies raised to complex I (CI) NADH-oxidoreductase subunit 9 (NAD9) and γ-carbonic anhydrase-like subunit 2 (CA2) proteins, Rieske iron-sulfur protein (RISP) of complex III (CIII), the cytochrome oxidase subunit 2 (COX2) of complex IV (CIV), mitochondrial ATPb subunit of ATP-synthase (CV) (Supplemental Table S3), as indicated in each blot. *In-gel* complex I activity assays were performed essentially as described previously (Meyer, *et al.* 2009). Arrows indicate to the native supercomplex I+III2 (~1,500 kDa), holo-CI (~1,000 kDa), III_2_ (~500 kDa), CIV (~220 kDa) and CV (~600 kDa), while I* indicates the presence of CI assembly intermediates.

BN-PAGE followed by immunoblotting with anti-RISP antibodies indicated to reduced CIII_2_ levels in *nmat3-1* mutant-line (Fig. 7). We also noticed the appearance of a higher molecular particle of *ca*. 700 kDa in *nmat3-1* plants of a yet unknown protein composition. Reduced CIII_2_ levels and the appearance of the higher molecular mass may relate to altered assembly and biogenesis of CI-CIII ‘super-complexes’. Accordingly, the higher mass particle was not apparent in the functionally complemented *nmat3*:35S nMAT3 line. Notably, the COX native blots also indicated a strong reduction in holo-CIV, which was associated with the appearance of a higher molecular mass particle of *ca*. 500 kDa containing the COX protein (Fig. 7). The COX2-associated 500 kDa particle was also observed in rescued Col-0 plants, suggesting a tissue or developmental altered CIV assembly, albeit at a lower level. The levels of the 500 kDa particle in *nmat3-1* plants were equivalent to the levels of the holo-CI in rescued Col-0 plants, the heterozygous line and the *nmat3*:35S nMAT3 complemented line (Fig. 7). As no changes in the expression of organellar genes encoding COX 1-3 subunits of the Cyt.C biogenesis factors, the molecular basis of these effects remains unclear and need to be further investigated. Immunoblots with anti-ATPb antibodies, indicated that CV accumulates to higher levels in rescued *nmat3*-1 and Col-0 plants (about 2.1 and 1.8 folds higher, respectively). As the accumulation of holo-CV was not affected in heterozygous *nmat3-1* plants or the complemented *nmat3*:35S nMAT3 line (about 1.03 and 0.97-fold, respectively) (Fig 7, and Table S2), we speculate that CV is likely upregulated during early seedlings development or due to the methodology used for rescuing the *nmat3*-1 and Col-0 plants.

### Arabidopsis *nmat3-1* mutants display complex I-specific respiratory defects

To analyze whether the respiratory activity was affected in *nmat3-1* mutant, the oxygen-uptake rates of 100 mg seedlings, obtained from 3-week-old MS-grown wild-type and 3-month-old *nmat3-1* mutant plants were monitored in the dark with a Clark-type electrode (Fig. 7). The average O_2_-uptake rates of *nmat3-1* (58.90 ± 2.62 nmol O_2_ min^−1^ grFW^−1^) was notably lower than that of wild-type plants (176.48 ± 12.36 nmol O_2_ min^−1^ grFW^−1^). The respiratory functions in wild-type and *nmat3* mutant were further analyzed in the presence of the electron transport inhibitors rotenone (ROT, 50 μM), antimycin A (Anti-A, 10 μM) and potassium cyanide (KCN, 1 mM). Pre-incubation with rotenone (50 μM), a specific complex I inhibitor, had a strong effect on the respiration rates of wild-type plants (*i.e*. 51.55 ± 4.31 nmol O_2_ min^−1^ gr FW^−1^) (Fig. 8, +ROT), about 71% lower than the O_2_-uptake rates under the standard conditions (Fig. 8, Mock). As also seen in various other mutants affected in CI biogenesis or activity, *e.g.* (Cohen, *et al.* 2014, Falcon de Longevialle *et al.* 2007, Shevtsov *et al.* 2018), rotenone had a milder effect on the respiratory activities of *nmat3* plants (*i.e*. 46.54 ± 2.79 nmol O_2_ min^−1^ grFW^−1^), about 21% inhibition from the control conditions (Fig. 8). Likewise treating the plants with Anti-A, a specific complex III inhibitor, had a marked effect (*i.e.* about 67% inhibition) on the respiratory activities of wild-type plants, and a milder effect on the O_2_-uptake rates of *nmat3-1* mutant plants (*i.e.* about 42% inhibition) (Fig. 8). The data shown in Figure 8 further indicated that the mitochondrial electron transport inhibition by CN^−^ is more pronounced in *nmat3-1* (26% inhibition), and also had a strong effect on the O_2_-uptake rates of wild-type plants (78% inhibition). These data indicate that CI, but not CII or CIV, was strongly affected in *nmat3-1* mutant, and may coincide with the upregulation of alternative pathways of electron transport, via the rotenone-insensitive type II NAD(P)H dehydrogenases (NDs) and the alternative oxidases (AOXs), which can bypass CI and CIV, respectively (see Fig. 6b).

**Figure 8.**
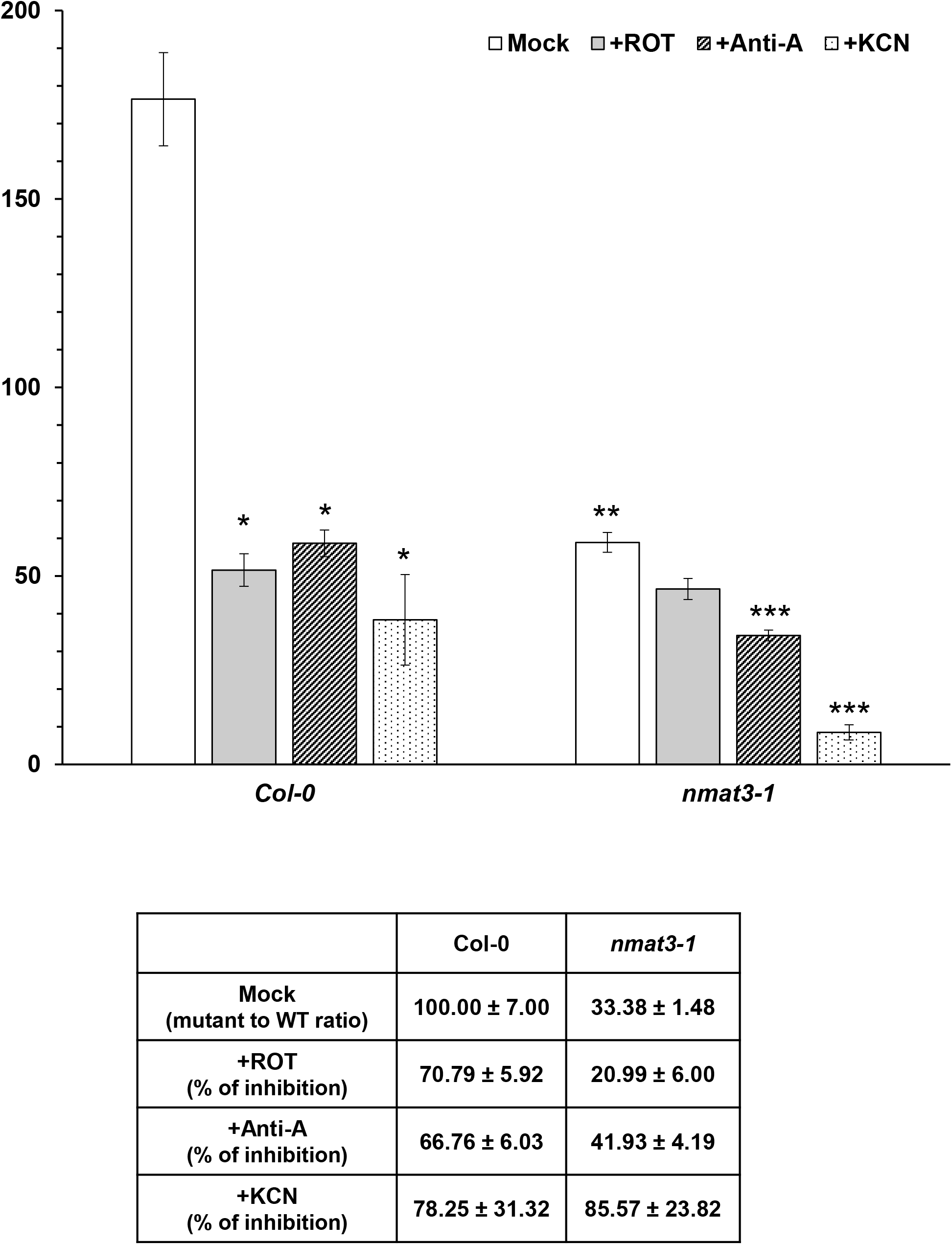
Respiration activity in wild-type and *nMAT3*-reduced lines. The O_2_-uptake rates of wild-type (Col-0) and embryo-rescued homozygous *nmat3-1* plants, were analyzed with a Clark-type electrode as described previously (Cohen, *et al.* 2014, Zmudjak, *et al.* 2017). For each assay, 100 mg MS-grown seedlings were submerged in 2.5 mL sterilized water and applied to the electrode in a sealed glass chamber in the dark. O_2_-uptake rates were measured in the absence (Mock) or presence of rotenone (ROT, 50 μM), Antimycin A (Anti-A, 10 μM) and KCN (1 mM) that inhibit complexes I, III and IV activities, respectively. The values are means of three biological replicates (error bars indicate one standard deviation), while asterisks indicate significant differences from wild-type plants (Student’s T-test, P 0.05).

## Discussion

### At5g04050 gene-locus encodes an essential mitochondria-localized maturase factor (nMAT3), required for embryo-development in Arabidopsis plants

The homology of nMAT3 with model maturases (Keren, *et al.* 2009, Mohr and Lambowitz 2003) and its localization by GFP-targeting studies (Keren, *et al.* 2009) strongly support a role for nMAT3 in the processing of mitochondrial group II introns in plants. The At5g04050 gene is predicted to encode two putative nMAT3 isoforms: *nMAT3.1* that encodes a 757 (i.e. corrected sequence) amino acid product that contains both the RT and D-En domains typical to group II intron-encoded endonucleases (Fig. 1b and Fig. S3), and a spliced variant (*nMAT3.2*), encoding a 694 amino acid long protein (Fig. 1b and Fig. S3). The postulated product of nMAT3.2 harbors a degenerated RT domain, lacking the highly conserved ‘RNA binding and splicing’ motif (*i.e.* the thumb or X domain) of group II maturases, and also lacks the C-terminal D-En domain (Fig. 1b and Fig. S3). Our data support the existence of only a single isoform, the *nMAT3.1* gene-product, in *Arabidopsis thaliana* (var. Col-0) plants (Fig. S3). Gene expression databases indicate that At5g04050 is expressed at low levels, with a dominant expression during early developmental stages (Fig. S4).

Macroscopic analyses of siliques obtained from the heterozygous *nmat3-1* (SAIL_254_F03) plants indicated a premature arrest of embryo development at the heart-torpedo transition stage (Fig. 2). Here, we sought to define the molecular functions of nMAT3, and to analyze its postulated roles in mtRNA metabolism by establishing loss-of-function mutations in the *nMAT3* gene-locus. As no homozygous mutant individuals could be identified among the mature seeds of self-fertilized heterozygous *nmat3-1* plants, we used a modified embryo-rescue approach used to recover germination-arrested phenotypes in Arabidopsis (Best, *et al.* 2019). The *in vitro* system allowed us to obtain homozygous *nmat3-1* plantlets, which displayed severe growth and developmental defect phenotypes (Fig. 2).

### nMAT3 is required for processing of pre-*nad1* transcripts in Arabidopsis mitochondria

Comparison between the mtRNA profiles of the homozygous *nmat3-1* mutant-line versus Arabidopsis wild-type plant indicated a strong perturbation in the maturation of *nad1* exons c-d, where the levels of *nad1* mRNA was found to be reduced by about 50,000-fold in the mutant (Fig. 4). Reduced levels (*i.e.* about 100-fold) were also apparent for *nad1* exons a-b, c-d and d-e. The accumulation of mature transcripts corresponding to other intron-containing genes, or to various ‘intronless’ transcripts, was similar to those of the wild-type plants. Analyses of the splicing efficiencies of the 23 introns in Arabidopsis mitochondria (Sloan *et al.* 2018, Unseld, *et al.* 1997), indicated a very strong defect in the *trans*-splicing of *nad1* i3 in rescued *nmat3-1*, where the splicing efficiency was reduced by ~20,000 fold in the mutant-line (Fig. 5). Altered splicing activities were also noted for *nad1* introns 1 and 4, where their splicing efficiencies in *nmat3-1* were found to be about 360 and 20-fold, respectively, lower than those of the wild-type plants (Fig. 5).

We also noted to reduced splicing efficiencies in the cases of *nad2* introns 1 and 2 (Fig. 5), but these were significantly less pronounced than those seen for *nad1* introns 3 and 4. The mild effects we see in *nad2* intron 1 and 2 splicing and the fact that the accumulation of their mRNAs were not affected by the mutation in *nMAT3* gene-locus suggest that the maturation of *nad2* is indirectly affected by the processing of *nad1* introns in Arabidopsis plants, as also previously noted for *otp43* (Falcon de Longevialle, *et al.* 2007), *nmat2* (Keren, *et al.* 2009) (Zmudjak, *et al.* 2017) and *nmat4* (Cohen, *et al.* 2014) mutants. The mechanisms by which altered expression of particular mt-RNAs influences global mitochondrial gene-expression or RNA metabolism are still under investigation.

We suggest that the altered mtRNA metabolism in *nmat3* is directly associated with the mutation in At5g04050 gene-locus. This view is strongly supported by the genetic and biochemical analyses. The mtRNA profiles of wild-type embryos rescued at the same developmental stage (i.e. late heart-to-torpedo) were equivalent to those of wild-type, whereas *nad1* was dramatically reduced in *nmat3-1* mutant. Moreover, functional complementation of *nmat3-1* restored the organellar RNA and protein profiles associated with the mutation in At5g04050 gene-locus (Figs. 4-7, *nmat3*:35S-nMAT3 line). Notably, while the mRNA levels of *nmat3*:35S-nMAT3 were equivalent to those of 3-week-old MS-grown and embryo-rescued wild-type plants (Fig. 4), the splicing efficiencies of many of organellar group II intron were increased in the complemented line (Fig. 5). We speculate that these effects may correspond to off-target effects by nMAT3, which is over-expressed under the 35S promoter in *nmat3* mutant (Fig. S6), or might be indirectly associated with the functions of MatR. The organellar MatR factor is encoded within *nad1* i4 of the *nad1.3* transcript, which is strongly upregulated in *nmat3* plants (Fig. 5). MatR was shown to function in the splicing of various group II introns (Sultan, *et al.* 2016), many of which their splicing is upregulated in the complemented line. It remains possible, however, that altered mtDNA expression caused by nMAT3 over-expression is also influenced by a plethora of metabolic changes and developmental stimuli (Falcon de Longevialle, *et al.* 2007, Keren, *et al.* 2012, Zmudjak, *et al.* 2013).

### Mitochondrial complex I biogenesis defects in *nmat3* mutants

Respiratory complex I (NADH:ubiquinone oxidoreductase, EC 7.1.1.2) is an L-shaped enzyme (Baradaran *et al.* 2013) that is embedded in the inner-mitochondrial membrane, and mediates the transfer of electrons from NADH to ubiquinone. The overall structure of CI has remained conserved from bacteria to animals and plants (Baradaran, *et al.* 2013). The biogenesis of the 1.0 MDa holo-CI in angiosperms involves the incorporation of ~50 different subunits, of both mitochondrial (NAD1, NAD2, NAD3, NAD4, NAD4L, NAD5, NAD6, NAD7 and NAD9) and nuclear origin (Lee *et al.* 2013). These are incorporated into two main assembly intermediates comprising the so-called ‘membrane’ and ‘peripheral’ arm-domains (Braun, *et al.* 2014, Klodmann *et al.* 2010, Ligas, *et al.* 2019, Ostersetzer-Biran 2016).

NAD1 is a central anchor component of mitochondrial complex I. In silico and cryo-EM analyses indicate that NAD1 is found in the core junction between the two arms of the native complex (Klodmann, *et al.* 2010, Ostersetzer-Biran 2016, Soufari *et al.* 2020). Likewise, genetic and biochemical data are indicating that a large reduction in NAD1 expression has a severe consequence on CI assembly and function (Ostersetzer-Biran 2016). The splicing defects in *nad1* we see in *nmat3* are expected to influence the biogenesis of holo-CI and the assembly of CI-associated supper complex. Immunoblots of crude organellar preparations indicate that while the CI subunits CA2 and NAD9 are apparent in *nmat3-1* mitochondria, NAD1 is below detectable levels in the mutant (Fig. 6). Accordingly, rescued *nmat3-1* seedlings show altered respiration and are less sensitive to inhibition by rotenone (Fig. 8), a specific CI inhibitor. BN-PAGE revealed that CI is below detectable levels in the *nmat3-1* mutant-line (Fig. 7). The biogenesis defects in CI are likely associated with reduced CI activity (Figs. 7 and 8) and the induction of the alternative electron pathways, as apparent in Figure 6b.

BN-PAGE followed by immunoblot assays with anti-CA2 antibodies further revealed the formation of partially assembled sub-CI intermediates of the membrane arm, of *ca.* 550, 500, 450 and 200 kDa in *nmat3-1* mitochondria (Fig. 7, labeled as I*). These results are in line with various other mutants that are affected in *nad1* expression, i.e. *nmat4* (Cohen, *et al.* 2014), *msp1025* (Best, *et al.* 2019) or *otp43* (Ligas, *et al.* 2019), which also accumulate CI assembly intermediates that lack complex I activity (Fig. 7). It should be noted, however, that our data are different from those of *opt43* and *nmat4* mutants that accumulate CI intermediates of *ca.* 200 kDa and 400 kDa, but not the 450 kDa or higher mass sub-CI particles (Fig. 7). Yet, similarly to *nmat3*, the *msp1025* mutant also accumulates higher mass sub-CI particles (Best, *et al.* 2019). The basis for these differences remains unknown, and may correspond to the levels of NAD1 in each mutant line. We speculate that NAD1 in plants might be incorporated into CI later during the membrane-arm assembly, or that CI can be assembled into a large non-functional high-molecular weight intermediates that lack NAD1 but contain CA2, as indicated by the immunoblots and in-gel activity assays (Fig. 7).

The signals of CIII_2_ and CIV were notably reduced in the mutant, and were accompanied by the appearances of higher mass particles that likely correspond to altered CIII and CIV complexes (Fig. 7). As no effects on the expression of organellar genes encoding CIII and CIV subunits, including COX1, COX2, COX3 and the cytochrome c biogenesis (CCM) factors, were observed in *nmat3-1* mutant, the molecular basis for these effects in the *nmat3* mutant-line remain unclear. It remains possible, however, that these relate to altered nuclear-gene expression, involving CIII and CIV biogenesis related-factors, or represent yet unknown pleiotropic effects that are in particularly ample during early plant development. These speculations await further investigation, although CI defects in plants were previously associated with delayed growth and altered cellular metabolism, as well as various secondary effects on photosynthetic activities (Dutilleul *et al.* 2003, Meyer *et al.* 2009, Priault *et al.* 2006). Various mutants affected in CI biogenesis show defective embryogenesis and low germination phenotypes, but are able to be maintained *in vitro* in media supplemented with vitamins and sugars (Ostersetzer-Biran 2016). The *wrinkled* morphology of the seeds and their dependency on external sources of energy indicate that seed maturation is incomplete in CI mutants, leading to the production of seeds with reduced reserves and viability (Keren, *et al.* 2012, Kuhn, *et al.* 2015). Recent data indicate that once photosynthesis has been established in the young seedlings, growth is no longer dependent on the application of external sugars.

### The maturation of *nad1* is a complex process, involving MatR and the four nuclear-encoded maturases

The expression of *nad1* in vascular plants is a complex process that may represent a key step in the regulation of complex I biogenesis (Ostersetzer-Biran 2016). Analyses of *nad1* mutants in *Chlamydomonas reinhardtii* and human mitochondria indicated that NAD1 incorporates at the very earliest stage of complex I assembly (Antonicka *et al.* 2003, Cardol *et al.* 2002). However, the incorporation of NAD1 and the molecular pathways that lead to CI assembly in land plants are still under investigation, and may be at variance with this view. At least 12 nuclear-encoded factors (Fig. 9) (Colas des Francs-Small and Small 2014) and one mitochondrial factor (*i.e*. MatR) (Sultan, *et al.* 2016), are required for the excision of the four introns within *nad1* in Arabidopsis mitochondria, and in each case involve at least one maturase protein. The different factors that function in *nad1* introns splicing are exemplified in Figure 9. These include the PPR factors MISF74 (Wang *et al.* 2018) (*nad1* i4) and OTP43 (Falcon de Longevialle, *et al.* 2007) (*nad1* i1), the RNA helicases ABO6 (He *et al.* 2012) (all four *nad1* introns) and PMH2 (Köhler *et al.* 2010, Zmudjak, *et al.* 2017) (*nad1* intron 2 and 3), the CRM-related mCSF1 protein (Zmudjak, *et al.* 2013) (*nad1* introns 2 and 3), a RAD52-like protein (Gualberto *et al.* 2015) (*nad1* i2), as well as the maturases-associated MatR protein (Sultan, *et al.* 2016) (*nad1* introns 3 and 4), nMAT1 (Keren, *et al.* 2012, Nakagawa and Sakurai 2006) (*nad1* i1), nMAT2 (Keren, *et al.* 2009, Zmudjak, *et al.* 2017) (*nad1* intron 2 and 3), nMAT3 (*nad1* introns 1, 3 and 4, this study) and nMAT4 (Cohen, *et al.* 2014) (*nad1* introns 1, 3 and 4). Remarkably, nMAT3 and nMAT4, which function in the splicing of the same subset of group II introns in Arabidopsis mitochondria, also seem to be evolutionarily related to one another (Brown, *et al.* 2014).

**Figure 9.**
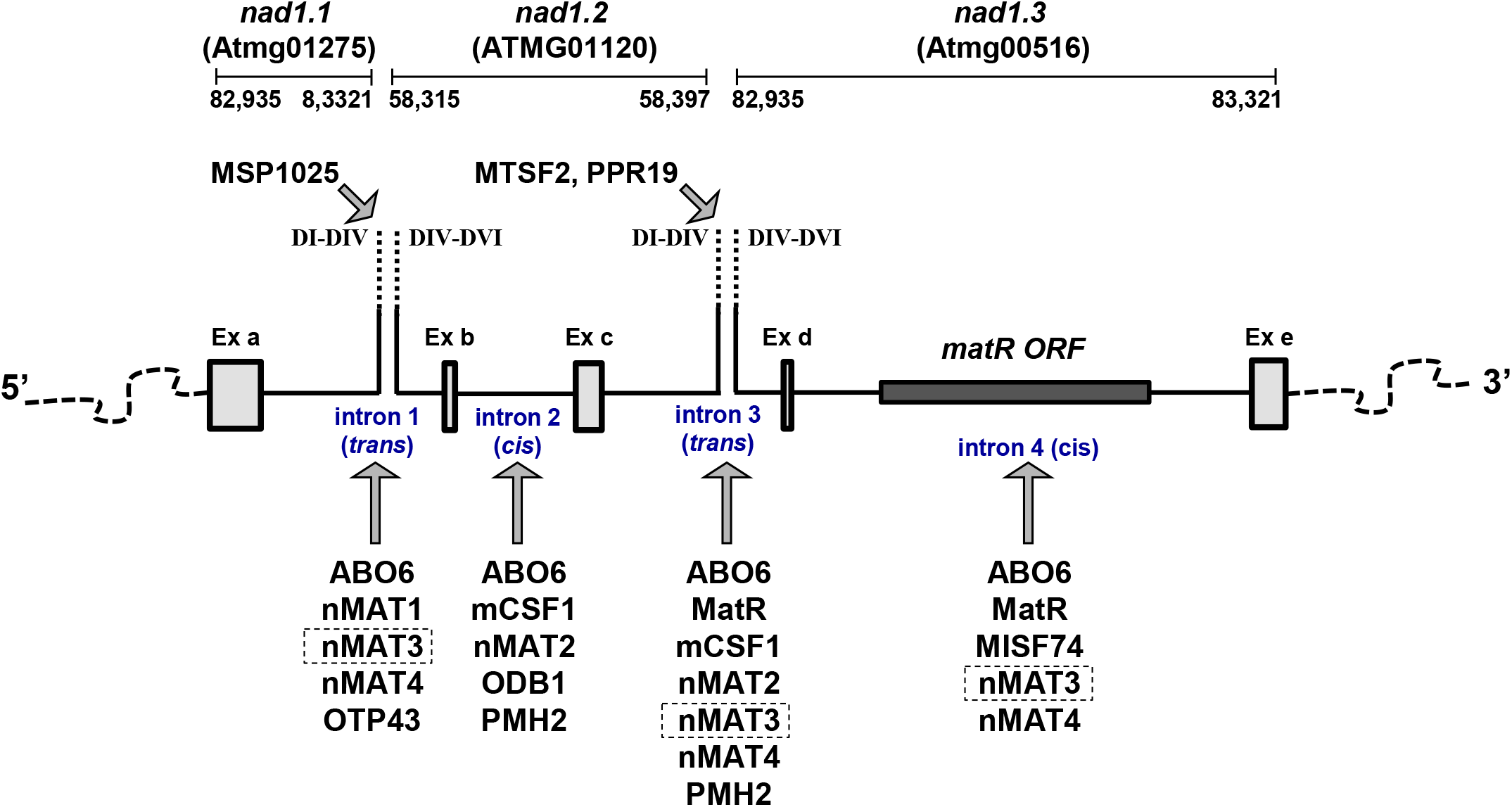
Target *nad1* introns and their accessory splicing cofactors. The maturation of *nad1* in Arabidopsis mitochondria involves the joining of five exons, encoded as three distinct transcription units, one spanning exon *nad1a* (i.e. *nad1.1*), a second spanning exons *nad1b* and *nad1c* (*nad1.1*) and a third spanning exons *nad1d* and *nad1e* (*nad1.1*), which are spliced through two *trans*- (introns 1 and 3) and two *cis*- (introns 2 and 4) events. The figure shows a schematic representation of *nad1* with exons (Ex) depicted as boxes and introns as lines. Specific gene loci are indicated according to the *de-novo* assembly of the Arabidopsis Col-0 mitogenome (BK010421) (Sloan, *et al.* 2018). The *trans*-spliced introns 1 and 3 of *nad1* are indicated as an interrupted dashed line. The *matR* reading frame is shown as a dark box within *nad1* intron 4. The figure represents various factors that have been demonstrated genetically to be involved for splicing of *nad1* introns. These include the PPR factors MISF74 (Wang, *et al.* 2018) and OTP43 (Falcon de Longevialle, *et al.* 2007), RNA helicases ABO6 (He, *et al.* 2012) and PMH2 (Köhler, *et al.* 2010, Zmudjak, *et al.* 2017), the CRM-related mCSF1 protein (Zmudjak, *et al.* 2013), the RAD52-like protein (Gualberto, *et al.* 2015), and the maturases MatR (Sultan, *et al.* 2016), nMAT1 (Keren, *et al.* 2012, Nakagawa and Sakurai 2006), nMAT2 (Keren, *et al.* 2009, Zmudjak, *et al.* 2017), nMAT3 (indicated by box) and nMAT4 (Cohen, *et al.* 2014). In addition to these factors, various PPR proteins were shown to determine cleavage sites required for the maturation and processing of group II introns, *e.g.* (Best, *et al.* 2019, Lee, *et al.* 2017, Wang, *et al.* 2017), or may recruit various essential splicing factors, as maturases and RNA helicases which are likely required to stabilize or to nucleate catalytically active structures required for splicing

The different group II intron splicing factors in Arabidopsis mitochondria are diverse in origin and most likely also in their mechanism of action. PPR proteins may affect intron structures by binding to their RNA targets in a sequence specific manner (Barkan and Small 2014), specify cleavage sites required for the intron maturation and processing. (*e.g.* (Best, *et al.* 2019, Lee *et al.* 2017, Wang *et al.* 2017), and may also recruit various essential splicing factors. Group II maturases and RNA helicases are likely required to stabilize or to nucleate catalytically active structures required for splicing (Brown, *et al.* 2014, Köhler, *et al.* 2010, Matsuura *et al.* 2001, Mohr *et al.* 2006, Ostersetzer *et al.* 2005, Schmitz-Linneweber, *et al.* 2015, Zmudjak, *et al.* 2017). The extreme complexity of NAD1 expression in Arabidopsis mitochondria further indicates its importance in regulating CI biogenesis in angiosperm species, and may also be associated with the nuclear control of respiratory functions during seed germination and early plant life (Best, *et al.* 2020). Moreover, the roles of the five mitochondria maturases in Arabidopsis, *e.g.* MatR, nMAT1, nMAT2, nMAT3 and nMAT4, in the splicing of many introns differ markedly from those of model maturases in bacteria and yeast mitochondria, which influence the processing of their own host introns or closely related RNAs (Schmitz-Linneweber, *et al.* 2015). This situation, which is also common to other mitochondrial splicing factors in plants, may represent a step in the gradual evolutionary transition from specific maturase-facilitated splicing of self-splicing introns towards the complex spliceosomal machinery in eukaryotes (Schmitz-Linneweber, *et al.* 2015). Characterization of group II intron splicing in plant mitochondria may also shed light on the functions and mechanisms of their proposed spliceosomal ‘descendants’ within the nucleus. The homology of the core splicing factor, Prp8, with maturases further supports this idea (Dlakic and Mushegian 2011).

### Physiological consequences of respiratory defects in *nmat3* mutants

Mitochondria have pivotal roles in aerobic cellular metabolism. Altered mitochondrial functions are often associated in plants with embryogenesis defects, reduced germination and growth or developmental-defect phenotypes. Embryo development and the ability to penetrate through the seed coat (Radchuk and Borisjuk 2014) require high amounts of metabolic energy that is provided through the metabolism of macromolecules stored in the seed (Best, *et al.* 2020). In some plants, as *Arabidopsis thaliana*, cotyledons absorb much of the endosperm nutrients and thus serve as the main energy supply source for the embryo. Accordingly, embryo development is strongly influenced by the size of the endosperm prior to its breakdown and absorbance by the cotyledons (Fourquin *et al.* 2016). Roughly on quarter of the seeds in heterozygous lines of *nma1*, *nmat4* and *otp43*, as well as other mutants affected in mitochondrial gene expression and RNA metabolism are shrunken and brown (Colas des Francs-Small and Small 2014). Similarly, about a quarter of the progeny in the siliques of *nmat3-1* were translucent seeds which contain embryos arrested at the heart to torpedo transition stage (Fig. 2).

Plant embryogenesis and seed-germination are complex developmental processes, which rely on cellular metabolism to support the high-energy demand for the developing embryos. Some Arabidopsis mutants with impaired embryogenesis at the cotyledon stages of development can be rescued on a nutrient medium designed to promote plant regeneration from immature wild-type cotyledons. Mutants arrested at earlier stages in embryogenesis can be also rescued, but these often fail to develop into mature seedlings (Franzmann, *et al.* 1989). The conditions for the embryo-rescue of *nmat3* mutants involved the addition of vitamins and sugars to promote complete embryo development, and hence seed-germination, and additionally required incubation in liquid medium to support the development beyond the L6 stage (Fig. 3). The growth and development of rescued *nmat3* plantlets was notably inhibited, as compare to rescued wild-type embryos of the same developmental stage (Fig. 3). These were not able to complete a life cycle, nor to flower or set seeds. Why does the loss of *nmat3* cause a major embryo defect, while the seeds of other homozygous mutants affected in *nad1* processing, *e.g.*, *nmat1*, *nmat2, nmat4* or *otp43* mutants, that also produce irregular wrinkled seed are viable? Currently, we cannot provide a definitive explanation to this phenomenon, but we speculate that the variations in growth and development between these mutants relate to the levels of NAD1. While *nmat1*, *nmat4* or *otp43* mutants seem to accumulate low levels of *nad1* mRNA, the level of NAD1 protein in *nmat3* is below detectable levels (Figs. 4-7). The accumulation of NAD1 in Arabidopsis may therefore relate with embryo maturation during seed development.

**In summary**, our genetic and biochemical data indicate that nMAT3 is a splicing factor that is pivotal for the expression of NAD1 in Arabidopsis mitochondria, cellular activities that are essential during early embryogenesis in Arabidopsis. Homozygous *nmat3* mutants are strongly affected in the splicing of *nad1 i3*, and also show reduced splicing efficiencies in *nad1* introns 1 and 4. Consequently, the levels of correctly processed *nad1* exons c-d RNA (and to a lesser degree also *nad1* exons a-b and d-e), are substantially reduced. The splicing defects associated with a marked decline in the accumulation of NAD1 protein and severe CI biogenesis defects, where holo-CI is below detectable levels in the *nmat3-1* mutant-line. The mtRNA profiles are related to the loss-of-function *nMAT3-1* allele, as rescued wild-type embryos at the late heart stage did not show any obvious effects on the accumulation or processing of *nad1* RNAs. Furthermore, the expression of *35S:nMAT3* in *nmat3-1* mutant restored the RNA metabolism defects and altered protein profiles seen in the homozygous line.

## Experimental procedures

### Plant material and growth conditions

*Arabidopsis thaliana* (ecotype Columbia) was used in all experiments. Wild-type (Col-0) and *nmat3* mutant lines, *i.e*. SAIL_254_F03 (*nmat3-1*), SALK_011307C (*nmat3-2*), SALK_139392 (*nmat3-3*) and SALK_144082 (*nmat3-4*) lines were obtained from ‘The Arabidopsis Information Resource’ (TAIR; http://www.arabidopsis.org) seed stock. Prior to germination, seeds obtained from wild-type and mutant lines were surface-sterilized with Cl_2_ gas, generated by the addition of 1.5 ml HCl per 50 mL of bleach (sodium hypochlorite 4.7%), for 4 hours at room-temperature (RT). The seeds were then sown on MS-agar plates containing 1% (w/v) sucrose or rescued by a method described in detail below. For synchronized germinations, the seeds were kept in the dark for 5 days at 4°C and then grown under long day condition (LD, 16:8-hour) in a controlled temperature and light growth chamber (Percival Scientific, Perry, IA, USA) at 22°C and light intensity of 300 μE m^−2^ s^−1^. PCR was used to screen the plant collection and check the insertion integrity of each individual line (specific oligonucleotides are listed in Supplemental Table S4). Sequencing of specific PCR products was used to analyze the precise insertion site in the T-DNA lines.

### Embryo-rescue and establishment of homozygous *nmat3* mutants

Green-matured siliques (i.e. about 10 days post anthesis), obtained from wild-type and heterozygous *nmat3-1* mutant plants, were surfaced sterilized with 6% bleach solution for 10 min at RT. The seeds were then soaked in a 70% ethanol solution for 10 min at RT, washed briefly with sterile DDW, and opened in a biological hood. Green (*i.e.*, wild-type or heterozygous) and white seeds (homozygous), were obtained from siliques of heterozygous *nMAT3-1*, at different developmental stages (i.e., distributed along the Inflorescence Stem). Likewise, immature seeds of Col-0 seeds at the heart stage were obtained from sterilized siliques. The seeds were then sown on MS-agar plates supplemented with 1% (w/v) sucrose and different vitamins (*i.e*., Myoinositol, Thiamine, Pyridoxine, and nicotinic acid). For DNA and RNA analysis we used Arabidopsis wild-type and *nMAT3* plantlets at stage L6 (*i.e.*, 6 to 8 leaves) (Boyes, *et al.* 2001). To obtain larger quantities of plant material, plantlets at stage L6 grown on MS-agar plates, were transplanted into a MS-based liquid medium supplemented with 1 to 3% (w/v) sucrose and vitamins (as above) and incubated in Arabidopsis growth chambers (Percival Scientific, Perry, IA, USA), under standard growth conditions with moderate (50~100 RPM) shaking (i.e., long day conditions, at 22°C and light intensity of 300 μE m^−2^ s^−1^).

### Plant transformation and functional complementation of the *nmat3-1* mutation

*nMAT3* clone was generated by subcloning of a 2,425 bp fragment containing the full coding region (including the “introns”) of *nMAT3* gene under the constitutive cauliflower mosaic virus 35S promoter (35S) or the seed-specific *ABI3* promoter (Despres, *et al.* 2001). The construct was cloned into pCAMBIA-2300 (Addgene). The shuttle vector containing the transgene was electroporated into Agrobacterium tumefaciens (strain GV3101), that was used to transform heterozygote *nmat3-1* plants by the floral dip method (Clough and Bent 1998). Seeds collected from individual T1 plants, were surface sterilized, and plated on MS-agar medium supplemented with kanamycin (50 μg/ml). Kanamycin-resistant T2 seedlings were transferred to the soil, grown to maturity and screened for the presence of the transgene (in the genetic background of homozygous *nmat3-1*). Seeds collected from positive plants were surface sterilized, plated on MS-agar medium supplemented with Kanamycin (50 μg/ml), and then analyzed for their RNA and protein profiles, organellar activities and plant phenotypes.

### Microscopic analyses of Arabidopsis wild-type and mutant plants

Analysis of whole plant morphology, roots, leaves, siliques and seeds of wild-type and mutant lines were examined under a Stereoscopic (dissecting) microscope or light microscope at the Bio-Imaging unit of the Institute of Life Sciences (The Hebrew University of Jerusalem).

### RNA extraction and analysis

RNA extraction and analysis were performed essentially as described previously (Cohen, *et al.* 2014, Keren *et al.* 2011, Shevtsov, *et al.* 2018, Sultan, *et al.* 2016, Zmudjak, *et al.* 2013). RNA was prepared from 50 mg seedlings grown on MS-agar plates supplemented with 1% sucrose, following standard TRIzol Reagent protocols (Ambion, Thermo Fisher Scientific) with additional phenol/chloroform extraction procedure. The RNA was treated with DNase I (RNase-free) (Ambion, Thermo Fisher Scientific) prior to its use in the assays. RT-qPCR was performed with specific oligonucleotides designed to amplify exon-exon (mRNAs) regions, corresponding to numerous mitochondrial genes, and intron-exon regions (pre-mRNAs) within each of the 23 group II introns in *Arabidopsis thaliana* (Table S1). Reverse transcription was carried out with the Superscript III reverse transcriptase (Invitrogen), using 3 - 5 μg of total RNA and 250 ng of a mixture of random hexanucleotides (Promega) and incubated for 50 min at 50°C. Reactions were stopped by 15 min incubation at 70°C and the RT samples served directly for real-time PCR. Quantitative PCR (qPCR) reactions were run on a LightCycler 480 (Roche), using 2.5 μL of LightCycler 480 SYBR Green I Master mix and 2.5 μM forward and reverse primers in a final volume of 5 μL. Reactions were performed in triplicate in the following conditions: pre-heating at 95°C for 10 min, followed by 40 cycles of 10 sec at 95°C, 10 sec at 58°C and 10 sec at 72°C. *GAPDH* (AT1G13440)*, ACTIN2* (At3g1878), *18S-rRNA* (At3g41768), *RRN26* (Atmg00020), *RRN5* (Atmg01380) and *RRN18* (Atmg01390), were used as reference genes in the qPCR analyses.

### Crude mitochondria preparation from MS-grown Arabidopsis seedlings

Crude mitochondria extracts were prepared essentially as described previously (Pineau *et al.* 2008, Shevtsov, *et al.* 2018). For the preparation of organellar extracts, about 200 mg of liquid MS-grown plantlets were harvested and homogenized in 2 ml of 75 mM MOPS-KOH, pH 7.6, 0.6 M sucrose, 4 mM EDTA, 0.2% polyvinylpyrrolidone-40, 8 mM L-cysteine, 0.2% bovine serum albumin and protease inhibitor cocktail ‘complete Mini’ from Roche Diagnostics GmbH (Mannheim, Germany). The lysate was filtrated through one layer of miracloth and centrifuged at 1,300 ×g for 4 min at 4°C (to remove cell debris). The supernatant was then centrifuged at 22,000 ×g for 10 min at 4°C. The resultant pellet, which contains thylakoid and mitochondrial membranes, was washed twice with 1ml of wash buffer 37.5 mM MOPS-KOH, 0.3 M sucrose and 2mM EDTA, pH 7.6.

### Protein extraction and analysis

An aliquot equivalent to 10 mg crude mitochondria extract or total protein extract were mixed with an equal volume of 3X protein sample buffer (Laemmli 1970), supplemented with 50 mM β-mercaptoethanol, and subjected to 12% SDS-PAGE (at a constant 100 V). Following electrophoresis, the proteins were transferred to a PVDF membrane (BioRad) and blotted overnight at 4°C with specific primary antibodies (Table S3). Detection was carried out by chemiluminescence assays after incubation with an appropriate horseradish peroxidase (HRP)-conjugated secondary antibody (Sigma or Santa Cruz).

### Blue native (BN) electrophoresis for isolation of native organellar complexes

Blue native (BN)-PAGE of crude organellar membranous fractions was performed generally according to the method described by Pineau, *et al.* (2008). An aliquot equivalent to 40 mg of crude Arabidopsis mitochondria extracts, obtained from wild-type and *nMAT3* plants was solubilized with 5% (w/v) digitonin in BN-solubilization buffer (30 mM HEPES, pH 7.4, 150 mM potassium acetate, 10% [v/v] glycerol), and then incubated on ice for 30 min. The samples were centrifuged 8 min at 20,000 xg to pellet any insoluble material and Serva Blue G (0.2% [v/v]) was added to the supernatant. The samples were then loaded onto a native 4 to 16% linear gradient gel. For ‘non-denaturing-PAGE’ immunoblotting, the gel was transferred to a PVDF membrane (Bio-Rad) in Cathode buffer (50 mM Tricine and 15 mM Bis-Tris-HCl, pH 7.0) for 16 h at 4°C at constant current of 40 mA. The blots where then incubated with antibodies (Table S3) against various organellar proteins, and detection was carried out by chemiluminescence assay with an appropriate horseradish peroxidase (HRP)-conjugated secondary antibody. For CI *in-gel* activity the gels were washed several times with DDW and the activity assays were performed essentially as described by Meyer, *et al.* (2009).

### Respiration activity

Oxygen consumption (*i.e.* O_2_ uptake) measurements were performed with a Clarke-type oxygen electrode, and the data feed was collected by Oxygraph-Plus version 1.01 software (Hansatech Instruments), as described previously (Shevtsov, *et al.* 2018). The electrode was calibrated with oxygen-saturated water and by the addition of excess sodium dithionite for complete depletion of the oxygen in water housed in the electrode chamber. Equal weights (200 mg) of wild-type and *NMAT3* seedlings were immersed in water and incubated in the dark for a period of 30 min. Total respiration was measured at 25°C in the dark following the addition of the seedlings to 2.5 mL of sterilized tap water in the presence or absence of rotenone (50 μM), Antimycin A (10 μM) and KCN (1 mM).

## Acknowledgments

We thank Dr. Michal Zmudjak for her help with establishing homozygous *nmat3* plants, and Nadav Biran-Ostersetzer for his help with figure preparation. We also wish to thank Dr. Etienne Meyer (Halle U.) for providing us with anti-NAD1 antibodies. The authors confirm that they have no conflict of interest to declare. This work was supported by grants to O.O.B from the ‘Israeli Science Foundation’ ISF grants no. 741/15 and 1834/20.

## Supplemental Materials

**Supplemental Figure S1. The nucleotide sequence of *nMAT3* gene and the locations of *nmat3-1* T-DNA-insertional line.**

The nucleotide sequence of *nMAT3*, encoded by At5g04050 gene-locus. Underlined letters indicate to the 5’ and 3’ untranslated regions (UTRs), as indicated by the RACE analysis and TAIR database, while uppercase letters represent the open reading frame of nMAT3. The position of T-DNA insertion in *nmat3-1* (SAIL_254_F03) line is indicated within the sequence of *nMAT3.* The location of the T-DNA insertion within nMAT3 gene was analyzed by PCR and sequencing. Red letters indicate to sequencing errors in ‘The Arabidopsis Biological Resource Center’ (ABRC) server that were confirmed by sequencing of the *nMAT3* gene-locus, as well as by BLAST searches against the Brassicales genome resources (see Supplemental Fig. S2).

**Supplemental Figure S2. Alignment of nMAT3 intron 1 sequence obtained by blast search of Brassicales genome sequences.**

Alignment was created using the NCBI nucleotide BLASTN server. Arabidopsis thaliana *nMAT3* gene (GenBank Accession At5g04050) confirmed the integrity of 3 independent sequencing data (The Hebrew University Genetic Resource). The aligned sequences were downloaded using the BLASTN server.

**Supplemental Figure S3. Analysis of the expression of the two-alternative spliced *nMAT3* isoforms in Arabidopsis thaliana (Col-0) plants.**

The At5g04050 gene-locus (Fig. S1) harbors two intron sequences that are suggested to be alternatively spliced into two isoforms: annotated as nMAT3.1, encoding a 757 amino acids protein, and the spliced variant, nMAT3.2, encoding a 694 amino acid protein product. The expression of the two putative gene products of At5g04050 was analyzed by RT-PCR with primers flanking introns 1 and 2 (Fig. S3). The unspliced product should yield a 751 nucleotides cDNA product, while the spliced isoform should yield a 500 nucleotide-long mRNA product. The RT-PCR data indicated the existence of only a single isoform, the unspliced *nMAT3.1* gene-product in Arabidopsis plants.

**Supplemental Figure S4. *nMAT3* gene expression patterns in different tissues and during various developmental stages.**

The expression patterns of nMAT3 were analyzed by publicly available microarray and high throughput sequencing databases, including (a) ‘The Arabidopsis Information Resource’ (TAIR; http://www.arabidopsis.org) and (b) Genevestigator analysis toolbox (Hruz, *et al.* 2008, Zimmermann, *et al.* 2004).

**Supplemental Figure S5. Accumulation of mitochondrial transcripts in different *nmat* mutants.**

Total RNA was extracted from 3-week-old seedlings of wild-type (Col-0) plants and *nmat1* (Keren, *et al.* 2012, Nakagawa and Sakurai 2006), *nmat2* (Keren, *et al.* 2009, Zmudjak, *et al.* 2017) and *nmat3* mutants. The relative transcript accumulation in the *nmat* mutants was analyzed by RNA gel blot hybridizations (i.e. northern blot analyses).

**Supplemental Figure S6. Accumulation of *nMAT3* transcripts in different *nmat3* mutants.**

RNA extracted from wild-type (Col-0), embryo-rescued *nmat3* mutants or Col-0 plants at the heart stage (Fig. 3) and *nmat3:35S-nMAT3* complemented plants was reverse-transcribed, and the relative steady-state levels of cDNAs corresponding to *nMAT3* were evaluated by qPCR with primers which specifically amplified the gene (Supplemental Table S4). The values are means of 9 RT-qPCR reactions corresponding to three biological replicates (error bars indicate one standard deviation), after normalization to the *ACTIN2* (At3g1878) and 18S-rRNA (At3g41768) genes.

**Supplemental Table S1.** Lists of oligonucleotides used for the analysis of the mRNA profiles of wild-type and *nmat3* mutant plants by RT-qPCR experiments.

**Supplemental Table S2.** Protein profiles quantification data.

**Supplemental Table S3.** List of antibodies used for the analysis of *nmat3* mutants.

**Supplemental Table S4.** Oligonucleotides used in screening of individual T-DNA insertion lines in Arabidopsis, and for the cloning of the *nMAT3* gene-fusion constructs.

## Author contributions statement

<u>Sofia Shevtsov-Tal</u>: Plant growth and analysis, embryo rescue and establishment of mutant lines, Biochemical analysis of gene expression, BN-PAGE analysis, RNA isolation, DNA sequencing, analysis of the transcriptome and splicing profiles of Arabidopsis mitochondria by RT-qPCR, respiration analyses. <u>Corinne Best</u>: assisted with RT-qPCR, immunoblots and the analyses of RNA and protein profiles. <u>Roei Matan</u>: assisting in DNA screening and RNA and protein analyses. <u>Dr. Sam Aldrin Chandran</u>: Plant growth and analyses, establishment of mutant lines. <u>Prof. Gregory G. Brown</u>: co-PI and manuscript preparation. <u>Prof. Oren Ostersetzer-Biran</u>: PI, manuscript preparation and corresponding author.

