## Supplemental for "nMAT3 is an essential maturase splicing factor required for holo-complex I biogenesis and embryo-development in *Arabidopsis thaliana* plants"

**_nMAT3_ gene structure**

aatttgttttctccctctctgctctgctcctcctccgtcgccgccggtgaacacttcgctgaacttttttctattccATGGTTTTGCGGCTGA
GAGTTCACTCTTTCTACAACCGTGGAATCTCATTCCTTGTCTCTTCTTCCTTGAGGAATCTTAGTACTGCTTCTTCGCTTTTTTTGAACTCC
GACCAAACCATCACTGAGCCATTGGTGAAATCAGAGCTCGAAGCCCTAGTTCTCAAGCAATACTCTCATGGTAAGTTCTACAGTCTCGTTAA
AAACGCCGTTTCTCTGCCTTGTGTCCTTCTCGCCGCCTGTCAGAATCTGTCTCTCTCCGCCAATTCGTCCGGAGATTTAGCCGATCGTGTCT
CCCGGAGGTTCTCAATCGAAGAAATGGGGCGCGAGATACGCGAAGGCAGGTTTGATATTCGTTCATGTTGCGTCGAATTCATCTCTTCAAGT
CTTGTTCTTCCCAATTTAAAACTCAAAGTGTTGATAGAAGCTATTAGAATGGTTCTTGAGATTGTGTATGATGATCGATTCGCTACGTTTTC
SAIL_254_F03
GTATGGTGGACGTGTTGGTATGGGTAGGCATACAGCGATTCGTTACCTGAAAA▼ACTCGGTTGAGAATCCACGGTGGTGGTTTCGTGTTTCG
TTTGCTCGAGAAATGTTTGAAGAACGTAATGTGGATATACTGTGTGGCTTTGTTGGAGAGAAGATCAACGATGTTATGTTGATTGAGATGAT
CAAGAAGCTGTTTGAGTTTGGGATTCTGAAGATTGAGCTTGGTGGATGTAATAGTGGAAGAGGTTTTCCACAAGAGTGTGGCTTGTGTTCGA
TTTTGATCAATGTTTACTTTGATGGACTTGACAAAGAGATTCAAGATTTGAGGCTAAAGATGAAAGTTAAGAATCCTCGCGTTGGTACAGGA
GATGAGGAATCAACAGGTAATGTTTTCTTTAAGCCAGTGAATATTTATGCAGTTAGGTATTTGGATGAGATACTTGTGATAACATCAGGCTC
GAAAATGCTGACGATGGATTTGAAGAAGCGGATTGTTGACATTTTGGAACAGAGACTAGAGTTGAGAGTAGATAGATTGAACACATCGATTC
ATAGTGCAGTGTCAGAGAAGATTAACTTTTTAGGGATGTATCTTCAGGCTGTTCCACCTTCGGTTTTGCGCCCGCCTAAGTCTGAGAAGGCG
GTCAGAGCAATGAAGAAGTATCAGAGGCAGAAAGACGTTAGGAAGTTGGAGTTAAGAAATGCTAGGGAGAGGAATCGTAAGACACTGGGGCT
TAAAATATTTAGACATGTCTTGAAGAAAATCAAGCAGAGCAATGGGTTTAAATTTGAAGGTGAGATAGAGAATGAGGTTAGGGATATTTTCC
AGAGCTGGGGAGAGGAAGTTATGCAAGACTTTATGGGTTCTCTCGAAGAGCGATGGAAATGGCACTGGCTGCTCACTAGAGGAGATTTTCTC
TCTTTGAGACATATTAGAGAGAAATTACCACAAGACCTGATTGATGCTTATGATGAATTTCAGGAGCAGGTGGACAAACATTTGGCTCCGAC
CCAGGCTAAAAAGGTACTTGAGGATGAGGAGAGGAGAGTAGAAGAAGAAGAGGAACAGAGATATGCTGAAAGGACAGTTGAGGATTTGACTA
AGTTATGCATGAAGGTGTCAGCTCCGGAGGAACTTGTCAGAAAGGCTATTAAgttagttgggttcacaaacagcatgggacgaccacggcct
atcattcacctcgtgactctcgaggattctgatatcatcaaatggtatgcaggggtagggcgtaaatggcttgatttcttctgctgttgcca
caattataagatggtaaaaatcattgttagttatcatatgaggttctcatgtattttgacactagcagagaagcatggatccactaaacgtg
aagctattagacactacacgaaagatctcagAGTATCAGATCTTGACGGGAGGGAAGAGGCCCATTTTCCATCGGAAAGAGAGGTTAAGATG
ATGGGAGACAAGAACCTTTCAGACCCTAAACCTGTGGATGGCACACTATCGTTGCTTTTGATCAGGCTCGCATCTGATGAGCCCTTGCATCA
CTGTGCTGCTAGTTTTTGTGAACGATCAGATACAATTATGCACCGTGTACATCTACTGCAAAACCGTTTACATATCAACCCACTAGACGAAG
AGAAATGGGTTCCTGGCATGGGCACCATCCACTCAGCTTTGAATCGGAAATGTCTCCCTTTGTGCTCCACTCACATCAGTGATGTATACTTG
GGTAAAATAACCCTTCAGGATGTTGATAGTAGTTCATTCATCGATCTCAGgtgatgtctgttactctcttctacaattggttataacatctg
cacgagagtaagaagtaaaaatggccattgctgagatcagGCGAAAATGCGTGCCAACAGGGTGTATCACATACACATTTTGTCTTGATTTC
ATCATGATAGGAAAATGAatgaatacttcataagcaactgtctggctgcgcaatggcatagtaagcttcataattgtcaggggggtttggttc
ttctcaggatcaggaaacaaagatttgagcattagttacctgaaaaacgcaatgatcatctgcatacacctgagctgtaccttgcacggagc
gaacatgtttgttcctttacggtcgtcagtatcacgctcagcattgtgatctcatttgacacaccaaaagcagttgacacgatggatatttg
tcgaattggcggttggcaattgacagttgtataactataattttgcaatcgtgtaatgtatgaatctacattttagtattgtagtagtaaga
gggacacttttcttctgtatcatgtaaagaaagaagtgaggtcaattactgttctggt

**Supplemental Figure S1. The nucleotide sequence of _nMAT3_ gene and the locations of _nmat3-1_ T-DNA-insertional line.**

The nucleotides sequence of *nMAT3*, encoded by At5g04050 gene-locus. Underlined letters indicate to the 5' and 3' untranslated regions (UTRs), as indicated by the RACE analysis and TAIR database, while uppercased letters represent the open reading frame of nMAT3. The position of T-DNA insertion in *nmat3-1* (SAIL_254_F03) line is indicated within the sequence of *nMAT3*. The location of the T-DNA insertion within nMAT3 gene was analyzed by PCR and sequencing. Red letters indicate to sequencing errors in 'The Arabidopsis Biological Resource Center' (ABRC) server that were confirmed by sequencing of the *nMAT3* gene-locus, as well as by BLAST searchers against the Brassicales genome resources (Supplemental Fig. S2).

**nMAT3 BLAST search results - Brassicales**

>LR782546.1:1102033-1102287 Arabidopsis thaliana genome assembly, chromosome: 5
GTTAGTTGGGTTCACAAACAGCATGGGACGACCACGGCCTATCATTCACCTTGTGACTCTCGAGGATTCTGATATCATCA
AATGGTATGCAGGGGTAGGGCGTAAATGGCTTGATTTCTTCTGCTGTTGCCACAATTATAAGATGGTAAAAATCATTGTT
AGTTATCATATGAGGTTCTCATGTATTTTGACACTAGCAGAGAAGCATGGATCCACTAAACGTGAAGCTATTAGACACTA
CACGAAAGATCTCAG
>LR699749.2:1082724-1082978 Arabidopsis thaliana genome assembly, chromosome: 5
GTTAGTTGGGTTCACAAACAGCATGGGACGACCACGGCCTATCATTCACCTTGTGACTCTCGAGGATTCTGATATCATCA
AATGGTATGCAGGGGTAGGGCGTAAATGGCTTGATTTCTTCTGCTGTTGCCACAATTATAAGATGGTAAAAATCATTGTT
AGTTATCATATGAGGTTCTCATGTATTTTGACACTAGCAGAGAAGCATGGATCCACTAAACGTGAAGCTATTAGACACTA
CACGAAAGATCTCAG
>LR699774.1:1093654-1093908 Arabidopsis thaliana genome assembly, chromosome: 5
GTTAGTTGGGTTCACAAACAGCATGGGACGACCACGGCCTATCATTCACCTTGTGACTCTCGAGGATTCTGATATCATCA
AATGGTATGCAGGGGTAGGGCGTAAATGGCTTGATTTCTTCTGCTGTTGCCACAATTATAAGATGGTAAAAATCATTGTT
AGTTATCATATGAGGTTCTCATGTATTTTGACACTAGCAGAGAAGCATGGATCCACTAAACGTGAAGCTATTAGACACTA
CACGAAAGATCTCAG
>LR699769.1:1116043-1116297 Arabidopsis thaliana genome assembly, chromosome: 5
GTTAGTTGGGTTCACAAACAGCATGGGACGACCACGGCCTATCATTCACCTTGTGACTCTCGAGGATTCTGATATCATCA
AATGGTATGCAGGGGTAGGGCGTAAATGGCTTGATTTCTTCTGCTGTTGCCACAATTATAAGATGGTAAAAATCATTGTT
AGTTATCATATGAGGTTCTCATGTATTTTGACACTAGCAGAGAAGCATGGATCCACTAAACGTGAAGCTATTAGACACTA
CACGAAAGATCTCAG
>LR699759.1:1085454-1085708 Arabidopsis thaliana genome assembly, chromosome: 5
GTTAGTTGGGTTCACAAACAGCATGGGACGACCACGGCCTATCATTCACCTTGTGACTCTCGAGGATTCTGATATCATCA
AATGGTATGCAGGGGTAGGGCGTAAATGGCTTGATTTCTTCTGCTGTTGCCACAATTATAAGATGGTAAAAATCATTGTT
AGTTATCATATGAGGTTCTCATGTATTTTGACACTAGCAGAGAAGCATGGATCCACTAAACGTGAAGCTATTAGACACTA
CACGAAAGATCTCAG
>LR699754.1:1103127-1103381 Arabidopsis thaliana genome assembly, chromosome: 5
GTTAGTTGGGTTCACAAACAGCATGGGACGACCACGGCCTATCATTCACCTTGTGACTCTCGAGGATTCTGATATCATCA
AATGGTATGCAGGGGTAGGGCGTAAATGGCTTGATTTCTTCTGCTGTTGCCACAATTATAAGATGGTAAAAATCATTGTT
AGTTATCATATGAGGTTCTCATGTATTTTGACACTAGCAGAGAAGCATGGATCCACTAAACGTGAAGCTATTAGACACTA
CACGAAAGATCTCAG
>LR215056.1:1093654-1093908 Arabidopsis thaliana genome assembly, chromosome: 5
GTTAGTTGGGTTCACAAACAGCATGGGACGACCACGGCCTATCATTCACCTTGTGACTCTCGAGGATTCTGATATCATCA
AATGGTATGCAGGGGTAGGGCGTAAATGGCTTGATTTCTTCTGCTGTTGCCACAATTATAAGATGGTAAAAATCATTGTT
AGTTATCATATGAGGTTCTCATGTATTTTGACACTAGCAGAGAAGCATGGATCCACTAAACGTGAAGCTATTAGACACTA
CACGAAAGATCTCAG
>LR881470.1:1100181-1100435 Arabidopsis thaliana genome assembly, chromosome: 5
GTTAGTTGGGTTCACAAACAGCATGGGACGACCACGGCCTATCATTCACCTTGTGACTCTCGAGGATTCTGATATCATCA
AATGGTATGCAGGGGTAGGGCGTAAATGGCTTGATTTCTTCTGCTGTTGCCACAATTATAAGATGGTAAAAATCATTGTT
AGTTATCATATGAGGTTCTCATGTATTTTGACACTAGCAGAGAAGCATGGATCCACTAAACGTGAAGCTATTAGACACTA
CACGAAAGATCTCAG
>LR797811.1:1091428-1091682 Arabidopsis thaliana genome assembly, chromosome: 5
GTTAGTTGGGTTCACAAACAGCATGGGACGACCACGGCCTATCATTCACCTTGTGACTCTCGAGGATTCTGATATCATCA
AATGGTATGCAGGGGTAGGGCGTAAATGGCTTGATTTCTTCTGCTGTTGCCACAATTATAAGATGGTAAAAATCATTGTT
AGTTATCATATGAGGTTCTCATGTATTTTGACACTAGCAGAGAAGCATGGATCCACTAAACGTGAAGCTATTAGACACTA
CACGAAAGATCTCAG
>LR797806.1:1120742-1120996 Arabidopsis thaliana genome assembly, chromosome: 5
GTTAGTTGGGTTCACAAACAGCATGGGACGACCACGGCCTATCATTCACCTTGTGACTCTCGAGGATTCTGATATCATCA
AATGGTATGCAGGGGTAGGGCGTAAATGGCTTGATTTCTTCTGCTGTTGCCACAATTATAAGATGGTAAAAATCATTGTT
AGTTATCATATGAGGTTCTCATGTATTTTGACACTAGCAGAGAAGCATGGATCCACTAAACGTGAAGCTATTAGACACTA
CACGAAAGATCTCAG
>LR797801.1:1140628-1140882 Arabidopsis thaliana genome assembly, chromosome: 5
GTTAGTTGGGTTCACAAACAGCATGGGACGACCACGGCCTATCATTCACCTTGTGACTCTCGAGGATTCTGATATCATCA
AATGGTATGCAGGGGTAGGGCGTAAATGGCTTGATTTCTTCTGCTGTTGCCACAATTATAAGATGGTAAAAATCATTGTT

AGTTATCATATGAGGTTCTCATGTATTTTGACACTAGCAGAGAAGCATGGATCCACTAAACGTGAAGCTATTAGACACTA
CACGAAAGATCTCAG
>LR797796.1:1098063-1098317 Arabidopsis thaliana genome assembly, chromosome: 5
GTTAGTTGGGTTCACAAACAGCATGGGACGACCACGGCCTATCATTCACCTTGTGACTCTCGAGGATTCTGATATCATCA
AATGGTATGCAGGGGTAGGGCGTAAATGGCTTGATTTCTTCTGCTGTTGCCACAATTATAAGATGGTAAAAATCATTGTT
AGTTATCATATGAGGTTCTCATGTATTTTGACACTAGCAGAGAAGCATGGATCCACTAAACGTGAAGCTATTAGACACTA
CACGAAAGATCTCAG
>AL391716.1:1266-1520 Arabidopsis thaliana DNA chromosome 5, BAC clone F21E1 (ESSA project)
GTTAGTTGGGTTCACAAACAGCATGGGACGACCACGGCCTATCATTCACCTTGTGACTCTCGAGGATTCTGATATCATCA
AATGGTATGCAGGGGTAGGGCGTAAATGGCTTGATTTCTTCTGCTGTTGCCACAATTATAAGATGGTAAAAATCATTGTT
AGTTATCATATGAGGTTCTCATGTATTTTGACACTAGCAGAGAAGCATGGATCCACTAAACGTGAAGCTATTAGACACTA
CACGAAAGATCTCAG
>LR699764.1:1120391-1120645 Arabidopsis thaliana genome assembly, chromosome: 5
GTTAGTTGGGTTCACAAACAGCATGGGACGACCACGGCCTATCATTCACCTTGTGACTCTCGAGGATTCTGATATCATCA
AATGGTATGCAGGGGTAGGGGGTAAATGGCTTGATTTCTTCTGCTGTTGCCACAATTATAAGATGGTAAAAATCATTGTT
AGTTATCATATGAGGTTCTCATGTATTTTGACACTAGCAGAGAAGCATGGATCCACTAAACGTGAAGCTATTAGACACTA
CACGAAAGATCTCAG
>LR797791.1:1122462-1122716 Arabidopsis thaliana genome assembly, chromosome: 5
GTTAGTTGGGTTCACAAACAGCATGGGACGACCACGGCCTATCATTCACCTTGTGACTCTCGAGGATTCTGATATCATCA
AATGGTATGCAGGGGTAGGGGGTAAATGGCTTGATTTCTTCTGCTGTTGCCACAATTATAAGATGGTAAAAATCATTGTT
AGTTATCATATGAGGTTCTCATGTATTTTGACACTAGCAGAGAAGCATGGATCCACTAAACGTGAAGCTATTAGACACTA
CACGAAAGATCTCAG
>AY094475.1:1708-1960 Arabidopsis thaliana AT5g04050/F8F6_260 mRNA sequence
GTTAGTTGGGTTCACAAACAGCATGGGACGACCACGGCCTATCATTCACCTTGTGACTCTCGAGGATTCTGATATCATCA
AATGGTATGCAGGGGTAGGGCGTAAATGGCTTGATTTCTTCTGCTGTTGCCAATTATAAGATGGTAAAAATCATTGTTAG
TTATCATATGAGGTTCTCATGTATTTTGACACTAGCAGAGAAGCATGGATCCACTAAACGTGAAGCTATTAGACACTACA
CGAAAGATCTCAG
>XM_021022764.1:1809-2063 PREDICTED: Arabidopsis lyrata subsp. lyrata uncharacterized LOC110227666 (LOC110227666), mRNA
GTTAGTTGGGTTCACAAACAACATGGGACGACCACGGCCTATCATTCACCTTGTGACTCTCGAGGATTCTGATATCATCA
AATGGTATGCGGGTGTAGGGCGTAAATGGCTTGATTTCTTCTGTTGTTGCCACAATTATAAGATGGTCAAAATCATTGTG
AGTTATCATATGAGGTTCTCATGTATTTTGACACTAGCAGAGAAGCACGGATCCACTAAACGTGAAGCTATTAGACACTA
CACGAAAGATCTCAG
>NM_120487.7:1708-1958 Arabidopsis thaliana RNA-directed DNA polymerase (reverse transcriptase) (AT5G04050), mRNA
GTTAGTTGGGTTCACAAACAGCATGGGACGACCACGGCCTATCATTCACCTCGTGACTCTCGAGGATTCTGATATCATCA
AATGGTATGCAGGGGTAGGGCGTAAATGGCTTGATTTCTTCTGCTGTTGCCACAATTATAAGATGGTAAAATCATTGGTA
CGTATCATATGAGGTTCTCATGTATTTGACACTAGCATGAGAAGCATGGATCCACTAAAAAGCTTATTAGACACTACACG
AAAGATCTCAG
>CP002688.1:1097723-1097973 Arabidopsis thaliana chromosome 5 sequence
GTTAGTTGGGTTCACAAACAGCATGGGACGACCACGGCCTATCATTCACCTCGTGACTCTCGAGGATTCTGATATCATCA
AATGGTATGCAGGGGTAGGGCGTAAATGGCTTGATTTCTTCTGCTGTTGCCACAATTATAAGATGGTAAAATCATTGGTA
CGTATCATATGAGGTTCTCATGTATTTGACACTAGCATGAGAAGCATGGATCCACTAAAAAGCTTATTAGACACTACACG
AAAGATCTCAG
>XM_010425337.2:1756-2010 PREDICTED: Camelina sativa uncharacterized LOC104708719 (LOC104708719), mRNA
GTTAGTTGGATTCACAAACAACATGGGACGGCCACGGCCTATCATTCACCTTACGAGTCTCGAGGATTCTGATATCATCA
AATGGTATGCGGGTATAGGGCGTAAATGGCTTGATTTCTTCTGCTGTTGCCACAATTATAAGATGGTCAAAATCATTGTA
AGTTATCATATGAGGTTTTCCTGTATTTTGACGCTAGCAGAGAAGCACGGATCCACTAAGCGTGAAGCTATTAGACACTA
CACGAAAGATCTCAG
>AL162873.1:86654-86906 Arabidopsis thaliana DNA chromosome 5, BAC clone F8F6 (ESSA project)
GTTAGTTGGGTTCACAAACAGCATGGGACGACCACGGCCTATCATTCACCTCGGTGACTCTCGAGGATTCTGATATCATC
AAATGGTATGCACGGGTAGGGCGTAAATGGCTTGATTTCTTCTGCTGTTGCCACAATTATAAGATCGGTAAAATCATTGG
TACGTATCATATGAGGTTCTCATGTATTTGACACTAGCATGAGAAGCATGGATCCACTAAAAAGCTTATTAGACACTACA
CGAAAGATCTCAG
>XM_010492598.2:1818-2072 PREDICTED: Camelina sativa uncharacterized LOC104768594 (LOC104768594), mRNA

GTTAGTTGGTTTCACAAACAGCATGGGACGGCCACGGCCTATCATTCACCTTACGAGTCTTGAGGATTCTGATATCATCA
AATGGTATGCGGGTATAGGGCGTAAATGGCTTGATTTTTTCTGCTGTTGCCACAATTATAAGATGGTCAAAATCATTGTA
AGTTATCATATGAGGTTTTCCTGTATTTTGACGCTAGCAGAGAAGCACGGATCCACTAAGCGTGAAGCTATTAGACACTA
CACGAAAGATCTCAG
>XM_019234965.1:1709-1963 PREDICTED: Camelina sativa uncharacterized LOC104734422 (LOC104734422), mRNA
GTTAGTGGGTTTCACAAACAGCATGGGACGGCCACGGCCTATCATTCACCTTACGAGTCTTGAGGATTCTGATATCATCA
AATGGTATGCGGGTATAGGGCGTAAATGGCTTGATTTCTTCTGCTGTTGCCACAATTATAAGATGGTCAAAATCATTGTA
AGTTATCATATGAGGTTTTCCTGTATTTTGACGCTAGCAGAAAAGCACGGATCCACTAAGCGTGAAGCTATTAGACACTA
CACGAAAGATCTCAG
>LR778317.1:32095224-32095476 Raphanus sativus genome assembly, chromosome: 8
TGAGATCTTTCGTGTAGTGTCTAATAGCTTCACGTTTAGTGGATCTGTGCTTCTCCGCTAGTGTCAAAATACAGGAGAAC
CTCATATGATAACTTACAATGATTTTGACCATCTTAAAGTTGTGGCAACAACAGAAGAAATCAAGCCATTTACGCCCTAC
ACCGGCATACCATTTGATGATATCAGAATCCTCAAGAGCCAAAAGGTGGCTGATAGGCCGTGGCCTTCCCATGCTGTTCG
TGAACCCAACTAA
>XM_018586784.1:1691-1943 PREDICTED: Raphanus sativus uncharacterized LOC108814249 (LOC108814249), mRNA
TTAGTTGGGTTCACAAACAACATGGGACGTCCACGGCCTATCAGCCACCTTTTGGCTCTTGAGGATTCTGATATCATCAA
ATGGTATGCGGGTGTAGGGCGTAAATGGCTTGACTTCTTCTGTTGTTGCCACAACTTTAAGATGGTCAAAATCATTGTAA
GTTATCATATGAGGTTCTCCTGTATTTTGACACTTGCAGAGAAGCACAGATCCACTAAACGTGAAGCTATTAGACACTAC
ACGAAAGATCTCA
>XM_024155928.1:1748-2000 PREDICTED: Eutrema salsugineum nuclear intron maturase 3, mitochondrial (LOC18016797), mRNA
TTAGTTGGCTTCACAAACAGCTTGGGCAGGCCACGGCCTATCAGCCACCTATTGGCTCTTGAGGATTCTGATATCATCAA
ATGGTATGCGGGTGTAGGGCGTAAATGGCTTGATTTCTTCTGTTGTTGCCACAATTATAGGATGGTCAAAATCATTGTAA
GTTATCACATGAGGTTCTCCTGTATTTTGACACTAGCAGAGAAGCACAGATCCACTAAACGTGAAGCTATTAGACACTAC
ACGAAAGATCTCA
>XM_023781755.1:1696-1950 PREDICTED: Capsella rubella nuclear intron maturase 3, mitochondrial (LOC111830203), mRNA
GTTAGTTGGTTTCACAAACAACATGGGACGACCACGGCCTATCGTTGACCTTACGTGTCTAGAAGATTCTGATATCATCA
AATGGTATGCGGGTATAGGGCGTAAATGGCTTGATTTCTTCTGCTGTTGCCACAATTATAAGATGCTCAAAATCATTGTA
AGTTATCATATGAGGTTTTCCTGTATTTTAACATTAGCAGAGAAGCACGGATCCACTAAACATGAAGCTATTAGACACTA
CACGAAAGATCTCAG
>LR031577.1:16939431-16939683 Brassica rapa genome, scaffold: A10
TGAGATCTTTCGTGTAGTGTCTAATAGCTTCACGTTTAGTGGATCTGTGCTTCTCCGCTAGTGTCAAAATACAGGAGAAT
CTCAGATGATAACTTACAATGATTTTGACCATCTTAAAGTTGTGGCAACAACAGAAGAAGTCAAGCCATTTACGCCCTAC
ACCCGCATACCATTTGATGATATCAGAATCCTCAAGAGCCAGAAGGTGGCTGATAGGCCGTGGCCGACCCATGTTGTTTG
TGAACCCAACTAA
>XM_013812932.2:1685-1937 PREDICTED: Brassica napus nuclear intron maturase 3, mitochondrial-like (LOC106372667), mRNA
TTAGTCGGGTTCACAAACAACATGGGACGGCCACGGCCTATCAGCCACCTTCTGGCTCTTGAGGATTCTGATATCATCAA
ATGGTATGCGGGTGTAGGGCGTAAATGGCTTGACTTCTTCTGTTGTTGCCACAACTTTAAGATGGTCAAAATCATTGTAA
GTTATCATATGAGGTTCTCCTGTATTTTGACTCTAGCGGAGAAGCACAGATCCACCAAACGTGAAGCTATTAGACACTAC
ACGAAAGATCTCA
>XM_018586785.1:1661-1913 PREDICTED: Raphanus sativus uncharacterized LOC108814251 (LOC108814251), mRNA
TTAGTTGGGTTCACGAACAGCATGGGACGTCCACGGCCCATCAGCCACCTTTTGGCTCTTGAGGATTCTGATATCATCAA
ATGGTATGCGGGTGTAGGGCGTAAATGGCTTGACTTCTTCTGTTGTTGCCACAACTTTAAGATGGTCAAAATCATTGTAA
GTTATCATATGAGGTTCTCCTGTATTTTGACACTTGCAGAGAAGCACAGATCCAGTAAACGCGAAGCTATTAGACACTAC
ACGAAAGATCTCA
>XM_009124392.3:2354-2606 PREDICTED: Brassica rapa nuclear intron maturase 3, mitochondrial (LOC103847334), mRNA
TTAGTCGGGTTCACAAACAACATGGGACGGCCACGGCCTATCAGCCACCTTCTGGCTCTTGAGGATTCTGATATCATCAA
ATGGTATGCGGGTGTAGGGCGTAAATGGCTTGACTTCTTCTGTTGTTGCCACAACTATAAGATGGTCAAAATCATTGTAA
GTTATCATCTGAGATTCTCCTGTATTTTGACTCTAGCGGAGAAGCACAGATCCACCAAACGTGAAGCTATTAGACACTAC
ACGAAAGATCTCA
>LR031875.1:62082289-62082541 Brassica oleracea HDEM genome, scaffold: C9
TGAGATCTTTCGTGTAGTGTCTAATAGCTTCACGTTTGGTGGATCTGTGCTTCTCCGCTAGTGTCAAAATACAGGAGAAT
CTCAGATGATAACTTACAATGATTTTCACCATCTTAAAGTTGTGGCAACAACAGAAGAAGTCAAGCCATGTACGCCCTAC
ACCCGCATACCATTTGATGATATCAGAATCCTCAAGAGCCAGAAGGTGGCTGATAGGCCGTGGCCGACCCATGTTGTTTG

TGAACCCAACTAA
>XM_013812741.2:1709-1961 PREDICTED: Brassica napus nuclear intron maturase 3, mitochondrial (LOC106372516), mRNA
TTAGTTGGGTTCACAAACAACATGGGTCGGCCACGGCCTATCAGCCACCTTCTGGCTCTTGAGGATTCTGATATCATCAA
ATGGTATGCGGGTGTAGGGCGTACATGGCTTGACTTCTTCTGTTGTTGCCACAACTTTAAGATGGTGAAAATCATTGTAA
GTTATCATCTGAGATTCTCCTGTATTTTGACACTAGCGGAGAAGCACAGATCCACCAAACGTGAAGCTATTAGACACTAC
ACGAAAGATCTCA
>LT669795.1:2140924-2141176 Arabis alpina genome assembly, chromosome: 8
TTAGTTGGGTTCACAAACAATATGGGTCGACCACGACCTATCAGCCACCTTCTGGTTCTTGAGGATTCTGATATCGTCAA
ATGGTATGCGGGAATAGGACGTAAATGGCTTGATTTCTTCTGTTGTTGCCACAATTATAAGATGGTCAAAATCATTGTAA
GTTATCATATGAGGTTCTCCTGTATTTTGACACTAGCAGAAAAGCACAGATCCACTAAACGTGAAGCTATTAAACACTAC
ACAAAAGATCTCA
>XM_013751025.1:1701-1953 PREDICTED: Brassica oleracea var. oleracea uncharacterized LOC106313257 (LOC106313257), mRNA
TTAGTTGGGTTCACAAACAACATGGGTCGGCCACGGCCTATCAGCCACCTTCTGGCTCTTGAGGATTCTGATATCATCAA
ATGGTATGCGGGTGTAGGGCGTACATGGCTTGACTTCTTCTGTTGTTGCCACAACTTTAAGATGGTGAAAATCATTGTAA
GTTATCATCTGAGATTCTCCTGTATTTTGACACTAGCGGAGAAGCACAGATCCACCAAACGTGAAGCTATTAGACACTAC
ACGAAAGATCTCA
>XM_010554675.2:1771-2022 PREDICTED: Tarenaya hassleriana uncharacterized LOC104823214 (LOC104823214), transcript variant X3, mRNA
GTTAGTCGGGTTCACGAACAACATGGGCCGGCCACGGCCAATCAGCCATCTTGTCGTTCTTGAAGATTCCGAAATCATCA
AATGGTACGCGGGAATCGGGAGAAGATGGCTTGATTTCTTCTGTTGTTGCCACAATTTCAAGATGATGAAAATAATCGTA
AGCTACCACATGAGGTTCTCCTGTATTTTGACATTAGCAGAGAAACACGAATCGACTAAACGTGAAGCTATTAAGCACTA
CACAAAGGATCT
>XM_010554674.1:1770-2021 PREDICTED: Tarenaya hassleriana uncharacterized LOC104823214 (LOC104823214), transcript variant X2, mRNA
GTTAGTCGGGTTCACGAACAACATGGGCCGGCCACGGCCAATCAGCCATCTTGTCGTTCTTGAAGATTCCGAAATCATCA
AATGGTACGCGGGAATCGGGAGAAGATGGCTTGATTTCTTCTGTTGTTGCCACAATTTCAAGATGATGAAAATAATCGTA
AGCTACCACATGAGGTTCTCCTGTATTTTGACATTAGCAGAGAAACACGAATCGACTAAACGTGAAGCTATTAAGCACTA
CACAAAGGATCT
>XM_010554672.1:1770-2021 PREDICTED: Tarenaya hassleriana uncharacterized LOC104823214 (LOC104823214), transcript variant X1, mRNA
GTTAGTCGGGTTCACGAACAACATGGGCCGGCCACGGCCAATCAGCCATCTTGTCGTTCTTGAAGATTCCGAAATCATCA
AATGGTACGCGGGAATCGGGAGAAGATGGCTTGATTTCTTCTGTTGTTGCCACAATTTCAAGATGATGAAAATAATCGTA
AGCTACCACATGAGGTTCTCCTGTATTTTGACATTAGCAGAGAAACACGAATCGACTAAACGTGAAGCTATTAAGCACTA
CACAAAGGATCT
>XM_022033057.1:3090-3324 PREDICTED: Carica papaya nuclear intron maturase 3, mitochondrial (LOC110807799), transcript variant X4, mRNA
GTCGGGTTCACAAACAACATGGGTCGTCCAAGGCCTATCGGCATACTTTTTGCCCTGGAAGATACTGATATTATTAAATG
GTATGCAGGTATTGGAAGAAGATGGCTTGAGTTCTTTTGTTGCTGCCACAACTTCAAGATGGTCAAAATTATTGTAAGCT
ATCACCTGAGGTTCTCCTGTATTTTGACATTAGCAGAAAAACATGGATCTACAAAAAAAGAAGCCATTAGTCACT
>XM_022033056.1:3540-3774 PREDICTED: Carica papaya nuclear intron maturase 3, mitochondrial (LOC110807799), transcript variant X3, mRNA
GTCGGGTTCACAAACAACATGGGTCGTCCAAGGCCTATCGGCATACTTTTTGCCCTGGAAGATACTGATATTATTAAATG
GTATGCAGGTATTGGAAGAAGATGGCTTGAGTTCTTTTGTTGCTGCCACAACTTCAAGATGGTCAAAATTATTGTAAGCT
ATCACCTGAGGTTCTCCTGTATTTTGACATTAGCAGAAAAACATGGATCTACAAAAAAAGAAGCCATTAGTCACT
>XM_022033055.1:4039-4273 PREDICTED: Carica papaya nuclear intron maturase 3, mitochondrial (LOC110807799), transcript variant X2, mRNA
GTCGGGTTCACAAACAACATGGGTCGTCCAAGGCCTATCGGCATACTTTTTGCCCTGGAAGATACTGATATTATTAAATG
GTATGCAGGTATTGGAAGAAGATGGCTTGAGTTCTTTTGTTGCTGCCACAACTTCAAGATGGTCAAAATTATTGTAAGCT
ATCACCTGAGGTTCTCCTGTATTTTGACATTAGCAGAAAAACATGGATCTACAAAAAAAGAAGCCATTAGTCACT
>XM_022033054.1:2285-2519 PREDICTED: Carica papaya nuclear intron maturase 3, mitochondrial (LOC110807799), transcript variant X1, mRNA
GTCGGGTTCACAAACAACATGGGTCGTCCAAGGCCTATCGGCATACTTTTTGCCCTGGAAGATACTGATATTATTAAATG
GTATGCAGGTATTGGAAGAAGATGGCTTGAGTTCTTTTGTTGCTGCCACAACTTCAAGATGGTCAAAATTATTGTAAGCT

ATCACCTGAGGTTCTCCTGTATTTTGACATTAGCAGAAAAACATGGATCTACAAAAAAAGAAGCCATTAGTCACT

>LR031876.1:4686966-4687007 Brassica oleracea HDEM genome, scaffold: C7

TTAATTGGGTTCACAAACAACATTGGACGGCCACGGCCTATC

**Supplemental Figure S2. Alignment of nMAT3 intron 1 sequence obtained by blast search of Brassicales genome sequences.**

Alignment was created using the NCBI nucleotide BLASTN server. Arabidopsis thaliana *nMAT3* gene (GenBank Accession At5g04050) confirmed the integrity of 3 independent sequencing data (The Hebrew University Genetic Resource). The aligned sequences were downloaded using the BLASTN server.

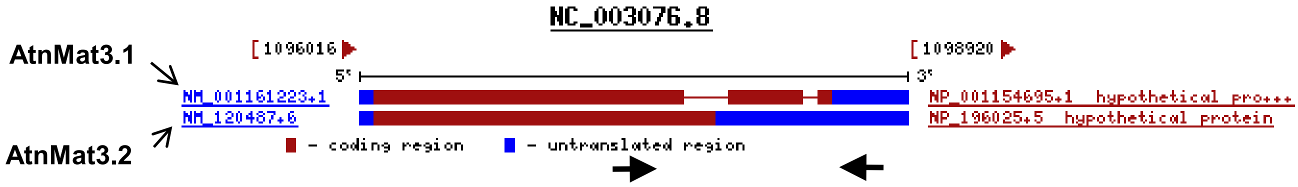


**Expected sizes:**

**At-nMat3.1 – 500 bp (spliced)**
**At-nMat3.2 – 751 bp (unspliced)**

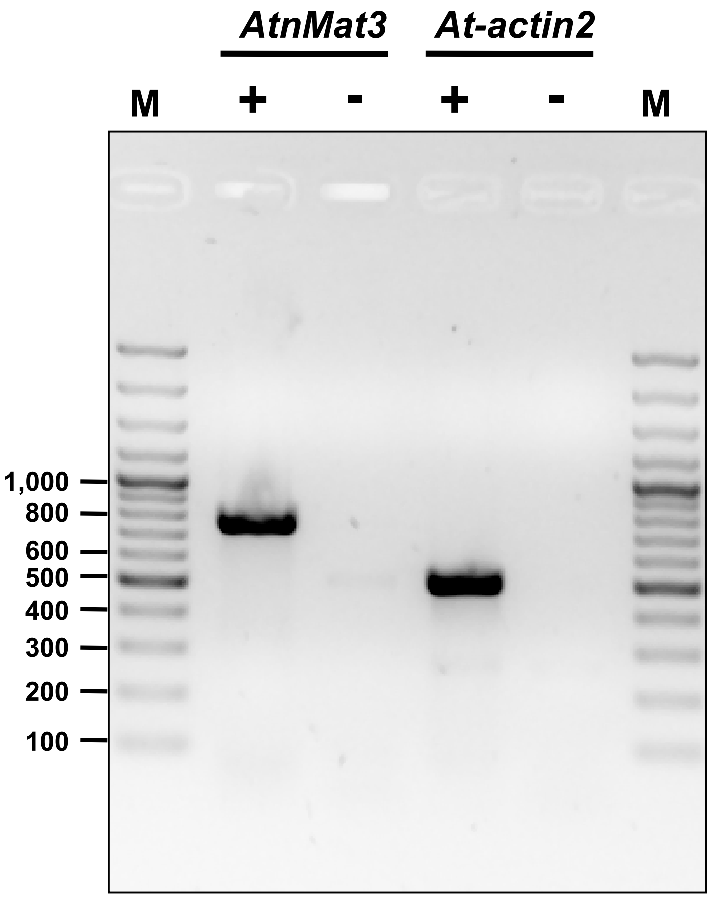


**Supplemental Figure S3. Analysis of the expression of the two-alternative spliced *nMAT3* isoforms in *Arabidopsis thaliana* (Col-0) plants.** The At5g04050 gene-locus (Fig. S1) harbors two intron sequences that are suggested to be alternatively spliced into two isoforms: annotated as nMAT3.1, encoding a 757 amino acids protein, and the spliced variant, nMAT3.2, encoding a 694 amino acid protein product. The expression of the two putative gene products of At5g04050 was analyzed by RT-PCR with primers flanking introns 1 and 2 (Fig. S3). The unspliced product should yield a 751 nucleotides cDNA product, while a spliced isoform should yield a 500 nucleotide-long mRNA product. The RT-PCR data indicated the existence of only a single isoform, the unspliced *nMAT3.1* gene-product in Arabidopsis plants.

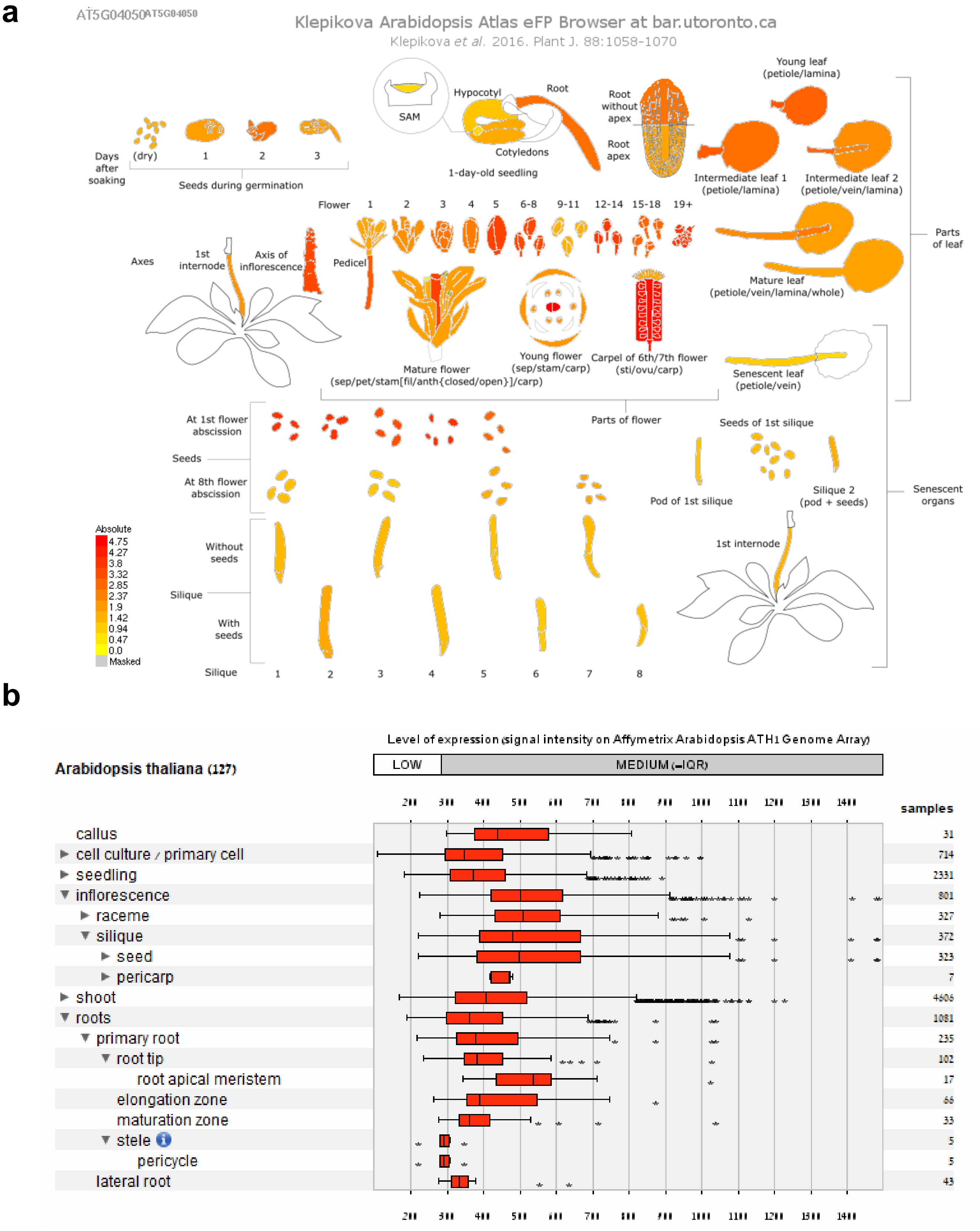


**Supplemental Figure S4.** ***nMAT3* gene expression patterns in different tissues and during various developmental stages.** The expression patterns of nMAT3 were analyzed by publicly available microarray and high throughput sequencing databases, including (a) 'The Arabidopsis Information Resource' (TAIR; http://www.arabidopsis.org) and (b) Genevestigator analysis toolbox (Hruz, *et al.* 2008, Zimmermann, *et al.* 2004).

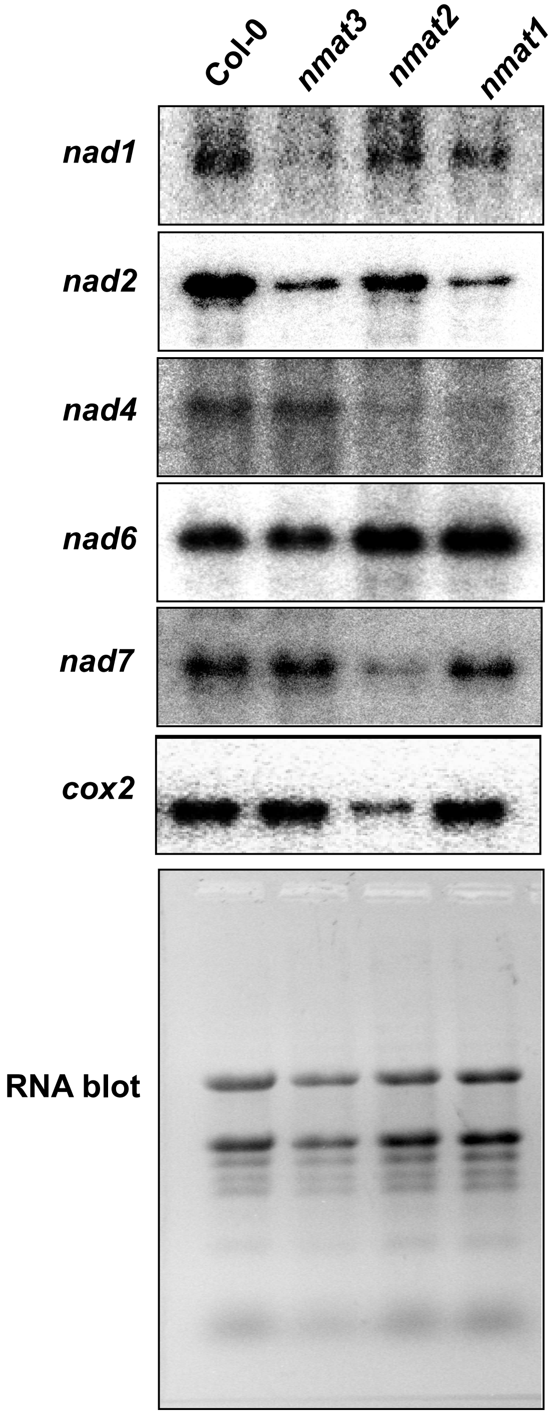


| | ***mutant to Col-0 ratio*** | | |
|---|---|---|---|
| gene | *nmat1* | *nmat2* | *nmat3* |
| *nad1* | 0.21 | 0.57 | 0.05 |
| *nad2* | 0.14 | 0.45 | 0.23 |
| *nad4* | 0.09 | 0.07 | 0.98 |
| *nad6* | 1.33 | 1.71 | 1.02 |
| *nad7* | 0.95 | 0.10 | 1.12 |
| *cox2* | 1.24 | 0.19 | 1.09 |

**Supplemental Figure S5. Accumulation of mitochondrial transcripts in different *nmat* mutants.** Total RNA was extracted from 3-week-old seedlings of wild-type (Col-0) plants and *nmat1* (Keren, *et al.* 2012a, Nakagawa and Sakurai 2006), *nmat2* (Keren, *et al.* 2009, Zmudjak, *et al.* 2017) and *nmat3* mutants. The relative transcript accumulation in the *nmat* mutants was analyzed by RNA gel blot hybridizations (i.e. northern blot analyses).

a

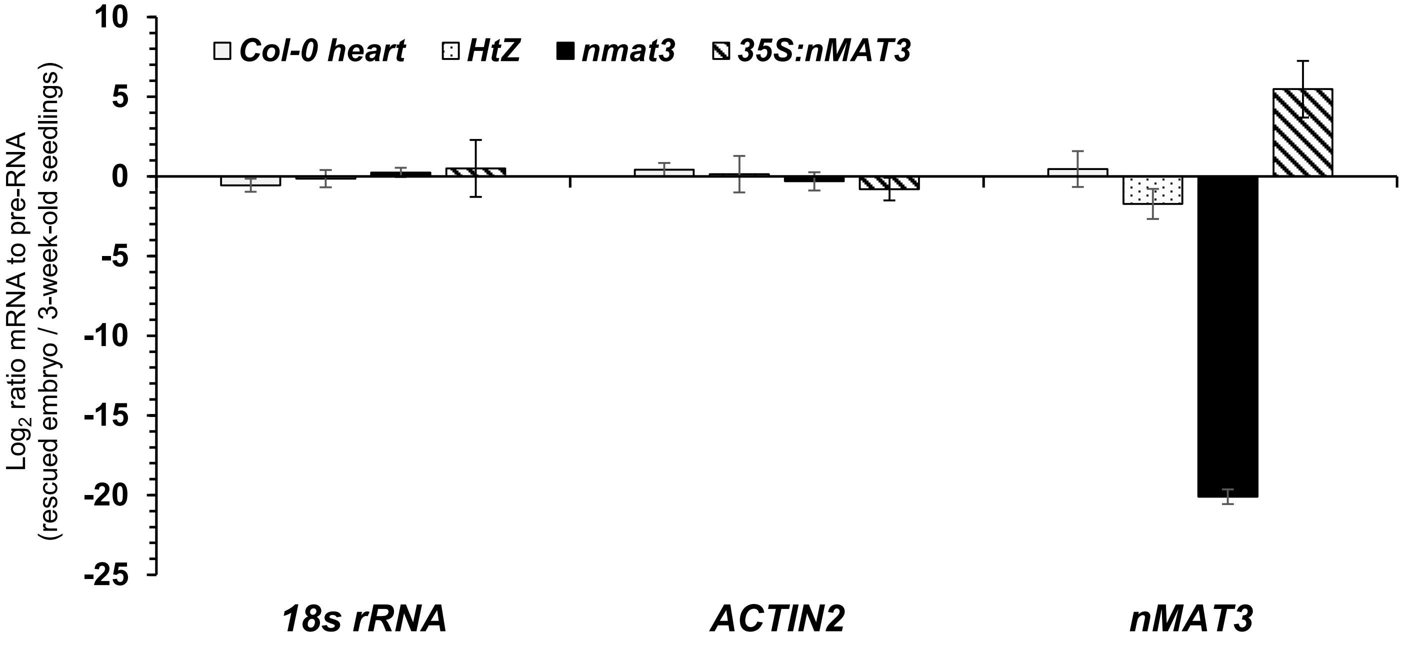


**Supplemental Figure S6. Accumulation of *nMAT3* transcripts in different *nmat3* mutants.**

RNA extracted from wild-type (Col-0), embryo-rescued *nmat3* mutants or Col-0 plants at the heart stage (Fig. 3) and *nmat3:35S-nMAT3* complemented plants was reverse-transcribed, and the relative steady-state levels of cDNAs corresponding to *nMAT3* were evaluated by qPCR with primers which specifically amplified the gene (Supplemental Table S4 ). The values are means of 9 RT-qPCR reactions corresponding to three biological replicates (error bars indicate one standard deviation), after normalization to the ACTINE2 (At3g1878) and 18S-rRNA (At3g41768) genes.

**Supplemental Table S1.**

a. Lists of oligonucleotides used for the analysis of the mRNA profiles of wild-type and mutant plants by RT-qPCR experiments.

| gene target | oligo name | sequence (5'-to-3') |
|---|---|---|
| *AOX1A* | *aox1aF* | AGCATCATGTTCCAACGACGTTTC |
| | *aox1aR* | GCTCGACATCCATATCTCCTCTGG |
| *AOX1B* | *aox1bF* | GGACCGTGAAATCTCTTCGATGGC |
| | *aox1bR* | TCTAGCATCATTGCTCTGCATCCG |
| *AOX1C* | *aox1cF* | TCTTCCAGAGGAGGTATGGTTGCC |
| | *aox1cR* | AGTGCATAAGCATCCCTCCAACC |
| *AOX1D* | *aox1dF* | TTTGCTCGAAGAGGCTGAGAACG |
| | *aox1dR* | CTCGTTCGTACCATTTGGGTTGTG |
| *AOX2* | *aox2F* | ACGGTGATTCGTGCTGATGAAGC |
| | *aox2R* | TCCTTGATTGCGAATGTCAGAAGC |
| *NDA1* | *nda1F* | GTATCCAACCGGCGATTTCACG |
| | *nda1R* | AGTTACAGTCTCACAATGCACCTC |
| *NDA2* | *nda2F* | TGGTGTTGGTCCTTCTCCTTTCG |
| | *nda2R* | TCCATTCGTCAATGCCAATCCTTC |
| *NDB1* | *ndb1F* | TAACACATTTGGCACTCCTGGTG |
| | *ndb1R* | CTCTGTGCATCCTCTACTTCCTTG |
| *atp1* | *atp1F* | TCACTTCGACACGTCTTTGC |
| | *atp1R* | GGAATGGCCTTGAATCTTGA |
| *atp6* | *atp6-1F* | TCTTTTGCGAGTCAATGCAC |
| | *atp6-1R* | TCTCGCGTATCTCACATTGC |
| *atp8* | *atp8F* | CCGTCGACTTATTGGGAAAA |
| | *atp8R* | TTCCTTGGCCATGTACAACA |
| *atp9* | *atp9F* | CATTCCCTCTGACGTCGAAT |
| | *atp9R* | TCGTCGATTCTTACCCTCGT |
| *atp4* | *atp4F* | GGATCAGCTTGCGAATTTGT |
| | *atp4R* | GCAAATTGCTTCCCCACTAA |
| *ccmb* | *ccmBF* | TCTTGGAATCACATCCAGCA |
| | *ccmBR* | CGAGACCGAAATTGGAAAAA |
| *ccmc* | *ccmCF* | AGCTACGCGCAAATTCTCAT |
| | *ccmCR* | GCCGTGGCGATATAAACAAT |
| *ccmfc* | *ccmFcF* | CACATGGAGGAGTGTGCATC |
| | *ccmFcR* | GTGGGTCCATGTAAATGATCG |
| *ccmfn-1* | *ccmFN1F* | AGCTCTTGGCATTGCTTTGT |
| | *ccmFN1R* | AGTGCCACAATCCCATTCAT |
| *ccmfn-2* | *ccmFN2F* | CGTGTCGTTCGTAATGGAAA |
| | *ccmFN2R* | TGATAAGCCCACCAACTTCC |
| *cob* | *cobF* | TGCCGGAATGGTATTTCCTA |
| | *cobR* | GCCAAAAGCAACCAAAACAT |
| *cox1* | *cox1F* | GTAGCTGCGGTGAAGTAGGC |
| | *cox1R* | CTGCCTGGATTCGGTATCAT |

| | | |
|---|---|---|
| *cox2* | *cox2F* | TGATGCTGTACCTGGTCGTT |
| | *cox2R* | TGGGGGATTAATTGATTGGA |
| *cox3* | *cox3F* | CCGTAACTTGGGCTCATCAT |
| | *cox3R* | AAACCATGAAAGCCTGTTGC |
| *mttb* | *mttBF* | GGGGTCTTTCTTTGGAAACC |
| | *mttBR* | TCTCCCTCATTCCACTCGTC |
| *nad1 exons a-b* | *nad1 1-2F* | GACCAATAGATACTTCATAAGAGACCA |
| | *nad1 1-2R* | TTGCCATATCTTCGCTAGGTG |
| *nad1 exons b-c* | *nad1 2-3F* | ATTCAGCTTCCGCTTCTGG |
| | *nad1 2-3R* | TCTGCAGCTCAAATGGTCTC |
| *nad1 exons c-d* | *nad1 3-4F* | AAAAGAGCAGACCCCATTGA |
| | *nad1 3-4R* | TCCGTTTGATCTCCCAGAAG |
| *nad1 exons d-e* | *nad1 4-5F* | AGCCCGGGATCTTCTTGA |
| | *nad1 4-5R* | TCTTCAATGGGGTCTGCTC |
| *nad2 exons a-b* | *nad2 exons a-bF* | GCGAGCAGAAGCAAGGTTAT |
| | *nad2 exons a-bR* | GGATCCTCCCACACATGTTC |
| *nad2 exons b-c* | *nad2 exons b-cF* | AAAGGAACTGCAGTGATCTTGA |
| | *nad2 exons b-cR* | AATATTTGATCTTAGGTGCATTTTC |
| *nad2 exons c-d* | *nad2 exons c-dF* | GCGCAATAGAAAGGAATGCT |
| | *nad2 exons c-dR* | CTATGGGTCTACTGGAGCTACCC |
| *nad2 exons d-e* | *nad2 exons d-eF* | CAAAGGAGAGGGGTATAGCAA |
| | *nad2 exons d-eR* | TATTTGTTCTTCGCCGCTTT |
| *nad3* | *nad3F* | CGAATGTGGTTTCGATCCTT |
| | *nad3R* | GCACCCCTTTTCCATTCATA |
| *nad4 exons a-b* | *nad4 exons a-bF* | ATTCTATGTTTTTCCCGAAAGC |
| | *nad4 exons a-bR* | GAAAAACTGATATGCTGCCTTG |
| *nad4 exons b-c* | *nad4 exons b-cF* | AATACCCATGTTTCCCGAAG |
| | *nad4 exons b-cR* | TGCTACCTCCAATTCCCTGT |
| *nad4 exons c-d* | *nad4 exons c-dF* | TTCCTCCATAAATTCTCCGATT |
| | *nad4 exons c-dR* | TGAAATTTGCCATGTTGCAC |
| *nad4L* | *nad4L-F* | GGGGAATCCTCCTTAATAGACG |
| | *nad4L-R* | AACGAAAATGGCTAACCCAATA |
| *nad5 exons a-b* | *nad5 exons a-bF* | TGGACCAAGCTACTTATGGATG |
| | *nad5 exons a-bR* | CCATGGATCTCATCGGAAAT |
| *nad5 exons b-c* | *nad5 exons b-cF* | TACCTAAACCAATCATCATATC |
| | *nad5 exons b-cR* | CTGGCTCTCGGGAGTCTCTT |
| *nad5 exons c-d* | *nad5 exons c-dF* | AACTCGGATTCGGCAAGAA |
| | *nad5 exons c-dR* | GATATGATGATTGGTTTAGGTA |
| *nad5 exons d-e* | *nad5 exons d-eF* | AACATTGCAAAGGCATAATGA |
| | *nad5 exons d-eR* | GTTCCTGCGTTTCGGATATG |
| *nad6* | *nad6F* | TATGCCGGAAAGGTACGAAG |
| | *nad6R* | GTGAGTGGGTCAGTCGTCCT |
| *nad7 exons a-b* | *nad7 exons a-bF* | ACCTCAACATCCTGCTGCTC |
| | *nad7 exons a-bR* | AAGGTAAAGCTTGAAGATAAGTTTTGT |
| *nad7 exons b-c* | *nad7 exons b-cF* | GAGGGACTGAGAAATTAATAGAGTACA |
| | *nad7 exons b-cR* | TGGTACCTCGCAATTCAAAA |
| *nad7 exons c-d* | *nad7 exons c-dF* | ACTGTCACTGCACAGCAAGC |

| | | |
|---|---|---|
| | *nad7 exons c-dR* | CATTGCACAATGATCCGAAG |
| *nad7 exons d-e* | *nad7 exons d-eF* | GATCAAAGCCGATGATCGTAA |
| | *nad7 exons d-eR* | AGGTGCTTCAACTGCGGTAT |
| *nad9* | *nad9F* | GGATGACCCTCGAAACCATA |
| | *nad9R* | CACGCATTCGTGTACAAACC |
| *rpl2* | *rpl2F* | CCGAAGACGGATCAAGGTAA |
| | *rpl2R* | CGCAATTCATCACCATTTTG |
| *rpl5* | *rpl5F* | AAGGGGTTCGACAGGAAAGT |
| | *rpl5R* | CGTATTTCGACCGGAAAATC |
| *rpl16* | *rpl16F* | GAGCATTTGCCAAACTCACA |
| | *rpl16R* | CGGACACTTTCATCGTGCTA |
| *rps3* | *rps3F* | CCGATTTCGGTAAGACTTGG |
| | *rps3R* | AGCCGAAGGTGAGTCTCGTA |
| *rps4* | *rps4F* | ACCCATCACAGAGATGCACA |
| | *rps4R* | TCACACAAACCCTTCGATGA |
| *rps7* | *rps7F* | CTCGAACTGAACGCGATGTA |
| | *rps7R* | AAGCTGCTTCAAGGATCCAA |
| *rps12* | *rps12F* | AGCCAAAGTACGGTTGAGCA |
| | *rps12R* | TTTGGGTTTTTCTGCACCAT |
| *matR* | *matR-F* | AATTTTTGCGAGAGCTGGAA |
| | *matR-R* | TTGAACCCCGTCCTGTAGAC |
| *rrn18* | *rrn18F* | CGTCACCTGGGTCAAAAACT |
| | *rrn18R* | GCTTGAAAACCGAAGTGAGC |
| *rrn26* | *rrn26F* | GACGAGACTTTCGCCTTTTG |
| | *rrn26R* | CTTGGAGCGAATTGGATGAT |
| *rrn5* | *rrn5F* | CCGACCTCGATATGTGGAATCGTC |
| | *rrn5R* | TGGACCATGTCTCCCGAACAATC |
| *18S rRNA* (nuclear) | *18S nucl-F* | AAACGGCTACCACATCCAAG |
| | *18S nucl-R* | ACTCGAAAGAGCCCGGTATT |
| *actin2* (At3g18780, nuclear) | *actin2-F* | GGTAACATTGTGCTCAGTGGTGG |
| | *actin2-R* | AACGACCTTAATCTTCATGCTGC |
| *GAPDH* | *GAPDH-F* | TCTCGATCTCAATTTCGCAAAA |
| | *GAPDH-R* | CGAAACCGTTGATTCCGATTC |

b. Lists of oligonucleotides used for the analysis of the splicing profiles of wild-type and mutant plants by RT-qPCR experiments.

| **Gene** | **Forward primer** | **Reverse primer** |
|---|---|---|
| *rpl2* | CCGAAGACGGATCAAGGTAA | CGCAATTCATCACCATTTTG |
| *rpl2* intron exon2 | TTAGGAAGAGCCGTACGAGG | CGCAATTCATCACCATTTTG |
| *rps3* | AGCCGAAGGTGAGTCTCGTA | CCGATTTCGGTAAGACTTGG |
| *rps3* intron1 exon2 | AGCCGAAGGTGAGTCTCGTA | TCTACGGCGGGGTCACTAT |
| *cox2* | TGGGGGATTAATTGATTGGA | TGATGCTGTACCTGGTCGTT |
| *cox2* intron1 exon2 | TGGGGGATTAATTGATTGGA | AGCAGTACGAGCTGAAAGGC |
| *ccmFc* | GTGGGTCCATGTAAATGATCG | CACATGGAGGAGTGTGCATC |
| *ccmFc* intron1 exon1 | CCCGGATCGAATCAGAGTT | CACATGGAGGAGTGTGCATC |
| *nad1* exon1-2 | GACCAATAGATACTTCATAAGAGACCA | TTGCCATATCTTCGCTAGGTG |

| | | |
|---|---|---|
| *nad1* intron1 exon2 | GACCAATAGATACTTCATAAGAGACCA | CGTGCTCGTACGGTTCATAG |
| *nad1* exon2-3 | ATTCAGCTTCCGCTTCTGG | TCTGCAGCTCAAATGGTCTC |
| *nad1* intron2 exon2 | GGTTGGGTTAGGGGAACATC | TCTGCAGCTCAAATGGTCTC |
| *nad1* exon3-4 | AAAAGAGCAGACCCCATTGA | TCCGTTTGATCTCCCAGAAG |
| *nad1* intron3 exon4 | AAAAGAGCAGACCCCATTGA | GGGAGCTGTATGAGCGGTAA |
| *nad1* exon4-5 | AGCCCGGGATCTTCTTGA | TCTTCAATGGGGTCTGCTC |
| *nad1* intron4 exon5 | AGCCCGGGATCTTCTTGA | ACGGAGCTGCATCCCTACT |
| *nad2* exon1-2 | GCGAGCAGAAGCAAGGTTAT | GGATCCTCCCACACATGTTC |
| *nad2* intron1 exon2 | GCGAGCAGAAGCAAGGTTAT | CCCATTCCTAACCAGTGGAG |
| *nad2* exon2-3 | AAAGGAACTGCAGTGATCTTGA | AATATTTGATCTTAGGTGCATTTTC |
| *nad2* intron2 exon2 | CCCGATCCGATAGTTTACAA | AATATTTGATCTTAGGTGCATTTTC |
| *nad2* exon3-4 | GCGCAATAGAAAGGAATGCT | CTATGGGTCTACTGGAGCTACCC |
| *nad2* intron3 exon4 | GCGCAATAGAAAGGAATGCT | GGCGAATTTCAAACTTGTGG |
| *nad2* exon4-5 | CAAAGGAGAGGGGTATAGCAA | TATTTGTTCTTCGCCGCTTT |
| *nad2* intron4exon4 | CTTATTCGTGGCAACCTTCC | TATTTGTTCTTCGCCGCTTT |
| *nad4* exon1-2 | ATTCTATGTTTTTCCCGAAAGC | GAAAAACTGATATGCTGCCTTG |
| *nad4* intron1 exon2 | CCGTATGATGCGGAAGTCTC | GAAAAACTGATATGCTGCCTTG |
| *nad4* exon2-3 | AATACCCATGTTTCCCGAAG | TGCTACCTCCAATTCCCTGT |
| *nad4* intron2 exon3 | GCGGAACGACCAGAAAAATA | TGCTACCTCCAATTCCCTGT |
| *nad4* exon3-4 | TTCCTCCATAAATTCTCCGATT | TGAAATTTGCCATGTTGCAC |
| *nad4* intron3 exon4 | TCTAGCTTGGTTCGGAGAGC | TGAAATTTGCCATGTTGCAC |
| *nad5* exon1-2 | TGGACCAAGCTACTTATGGATG | CCATGGATCTCATCGGAAAT |
| *nad5* intron1 exon2 | TGGACCAAGCTACTTATGGATG | TTCGCAAATAGGTCCGACT |
| *nad5* exon2-3 | TACCTAAACCAATCATCATATC | CTGGCTCTCGGGAGTCTCTT |
| nad5 intron2-exon2 | GTACGATCGTGTCGGGTGA | CTGGCTCTCGGGAGTCTCTT |
| *nad5* exon3-4 | AACTCGGATTCGGCAAGAA | GATATGATGATTGGTTTAGGTA |
| nad5 intron3-exon4 | AACTCGGATTCGGCAAGAA | GCCGTGTAATAGGCGACCA |
| *nad5* exon4-5 | AACATTGCAAAGGCATAATGA | GTTCCTGCGTTTCGGATATG |
| *nad5* intron4 exon5 | AACATTGCAAAGGCATAATGA | CCTGTAAACCCCCATGATGT |
| *nad7* exon1-2 | ACCTCAACATCCTGCTGCTC | AAGGTAAAGCTTGAAGATAAGTTTTGT |
| *nad7* intron1 exon2 | ACGGTTTTTAGGGGGATCTG | AAGGTAAAGCTTGAAGATAAGTTTTGT |
| *nad7* exon2-3 | GAGGGACTGAGAAATTAATAGAGTACA | TGGTACCTCGCAATTCAAAA |
| *nad7* intron2 exon3 | AGTGGGAGAGCCGTGTTATG | TGGTACCTCGCAATTCAAAA |
| *nad7* exon3-4 | ACTGTCACTGCACAGCAAGC | CATTGCACAATGATCCGAAG |
| *nad7* intron3 exon4 | TAAAGTGAAGTGGTGGGCCT | CATTGCACAATGATCCGAAG |
| *nad7* exon4-5 | GATCAAAGCCGATGATCGTAA | AGGTGCTTCAACTGCGGTAT |
| *nad7* intron4 exon5 | CGGCCAAATGACTACAGGAT | AGGTGCTTCAACTGCGGTAT |

**Supplemental Table S2**. Protein profiles quantification data.

Relative accumulation of native organellar respiratory complexes in wild-type and *nmat3* plants.

| Genetic line | holo-CI | | $CIII_2$ | CIV | CV |
|---|---|---|---|---|---|
| | CA2 | NAD9 | | | |
| **Col-0** | 1.00 | 1.00 | 1.00 | 1.00 | 1.00*** |
| **Col-0 heart** | 1.31 | 0.93 | 0.87 | 0.51** | 1.78*** |
| ***nmat3 (HtZ)*** | 0.88 | 0.66 | 1.15 | 0.47 | 1.03*** |
| ***nmat3 (HmZ)*** | LoD* | LoD* | 0.37** | LoD** | 2.14 |
| **Comp. 35S:*NMAT3*** | 0.72 | 0.54 | 0.98 | 0.53 | 0.97*** |

* - LoD - below limit of detection
** - appearance of an additional protein band of higher mass
*** - appearance of an additional protein band of lower mass

**Supplemental Table S3.** List of antibodies used for the analysis of *nmat3* mutants.

| Antibody | Protein I.D. | origin | serum | dilution | Reference / source |
|---|---|---|---|---|---|
| CA2 | γ-carbonic anhydrase-like subunit 2 | *Arabidopsis thaliana* | Rabbit (polyclonal) | 1/1,000 | (Perales *et al.* 2005, Sunderhaus *et al.* 2006) |
| NAD1 | NADH-dehydrogenase complex subunit-1 | *Arabidopsis thaliana* | Rabbit (polyclonal) | 1/1000 | Gift of Dr. Etienne Meyer, Halle U. |
| NAD9 | NADH-dehydrogenase complex subunit-9 | *Triticum spp.* | Rabbit (polyclonal) | 1/50,000 | (Lamattina *et al.* 1993) |

**Supplemental Table S4.** Oligonucleotides used in screening of individual T-DNA insertion lines in Arabidopsis, and for the cloning of the *nMAT3* gene-fusion constructs.

| Gene target | Gene I.D. | oligo name | Oligonucleotide sequence (5'-to-3') |
|---|---|---|---|
| ***nMAT3 gene*** | At5g04050 | nMAT3 full F | ATGGTTTTGCGGCTGAGAGTT |
| | | nMAT3 full R | TCATTTTCCTATCATGATGAAATC |
| ***nMAT3 screening*** | At5g04050 | nMAT3 screening F | ATACGCGAAGGCAGGTTTGA |
| | | nMAT3 full R | TCATTTTCCTATCATGATGAAATC |
| ***nMAT3 gene* expression** | At5g04050 | Ex nmat3 set1 F | ATGGTGGACGTGTTGGTATG |
| | | Ex nmat3 set1 R | CTTCAAACATTTCTCGAGCAAA |
| **ACTIN2** | At3g18780 | Act2-F | TCTTCCGCTCTTTCTTTCCAAG |
| | | Act2-R | CTGGCGTACAAGGAGAGAAC |
| **SALK T-DNA right border** | --- | RB-10064 | CCAGATCCGGTGCAGATTATTTG |
| **SALK T-DNA left border** | --- | LB-6443 | CATCGCCCTGATAGACGGTT |
| **SALK T-DNA left border** | --- | LBb1.3 | ATTTTGCCGATTTCGGAAC |
| **SAIL T-DNA left border** | --- | Ftsh_LB | CCTATTATATCTTCCCAAATTACCAATACA |
